# Positional motif analysis reveals the extent of specificity of protein-RNA interactions observed by CLIP

**DOI:** 10.1101/2021.12.07.471544

**Authors:** Klara Kuret, Aram Gustav Amalietti, Jernej Ule

**Affiliations:** National Institute of Chemistry, Hajdrihova 19, SI-1001 Ljubljana, Slovenia; The Francis Crick Institute, 1 Midland Road, London NW1 1AT, UK; Department of Neuromuscular Diseases, UCL Queen Square Institute of Neurology, Queen Square, London, WC1N 3BG, UK; Jozef Stefan International Postgraduate School, Jamova cesta 39, 1000 Ljubljana, Slovenia

**Keywords:** RNA-binding protein, protein-RNA interaction, CLIP, k-mer, RNA binding specificity, RNA motif, low-complexity region

## Abstract

**Background:** Crosslinking and immunoprecipitation (CLIP) is a method used to identify *in vivo* RNA– protein binding sites on a transcriptome-wide scale. With the increasing amounts of available data for RNA-binding proteins (RBPs), it is important to understand to what degree the enriched motifs specify the RNA binding profiles of RBPs in cells.

**Results:** We develop positionally-enriched k-mer analysis (PEKA), a computational tool for efficient analysis of enriched motifs from individual CLIP datasets, which minimises the impact of technical and regional genomic biases by internal data normalisation. We cross-validate PEKA with mCross, and show that background correction by size-matched input doesn’t generally improve the specificity of detected motifs. We identify motif classes with common enrichment patterns across eCLIP datasets and across RNA regions, while also observing variations in the specificity and the extent of motif enrichment across eCLIP datasets, between variant CLIP protocols, and between CLIP and *in vitro* binding data. Thereby we gain insights into the contributions of technical and regional genomic biases to the enriched motifs, and find how motif enrichment features relate to the domain composition and low-complexity regions (LCRs) of the studied proteins.

**Conclusions:** Our study provides insights into the overall contributions of regional binding preferences, protein domains and LCRs to the specificity of protein-RNA interactions, and shows the value of cross-motif and cross-RBP comparison for data interpretation. Our results are presented for exploratory analysis via an online platform in an RBP-centric and motif-centric manner (https://imaps.goodwright.com/apps/peka/). PEKA is available from https://github.com/ulelab/peka.

## Background

RNA regulation is mediated through dynamic interactions with RNA-binding proteins (RBPs) [1]. To understand how RBP-RNA interactions direct cellular processes, it is first necessary to identify RBP binding sites on the RNA, which is commonly achieved with crosslinking and immunoprecipitation (CLIP) technologies [2]. To date, the largest resource of CLIP data has been produced using the eCLIP method on 150 RBPs by the ENCODE consortium [3–5], thus enabling systematic studies of the features that recruit RBPs to specific RNA sites. While these data have been used to identify the sequence motifs bound by RBPs, the extent of specificity and enrichment of these motifs has not yet been compared across RBPs. Such analysis could inform on the extent to which RNA sequence determines the specificity of protein-RNA interactions, as well as on the quality and specificity of the CLIP data itself.

CLIP uses UV light to crosslink the direct protein-RNA contacts, followed by recovery of crosslinked RNA fragments to obtain a transcriptome-wide crosslinking landscape of a specific protein [6]. After crosslinking, the cells are lysed, RNA is partially fragmented, and the selected RBP is purified along with the crosslinked RNA fragments, which are used to prepare cDNA libraries for high-throughput sequencing. Resulting sequencing data is processed initially to identify the genomic positions of the nucleotides that crosslinked to the studied RBP and then leverage these positions to obtain regions of high RBP crosslinking, termed peaks. Subsequently, peaks are used to derive features driving the specificity of RBP binding, such as RNA sequence motifs or RNA structural elements.

Many methods have been developed for motif discovery from CLIP data [7]. Some of the more recent approaches use machine learning for comparative analyses of dozens of RBPs at once, but the applicability of this approach depends on availability of large CLIP resources with low technical variations or batch effects between datasets [8–13]. Conversely, algorithms that identify biologically-relevant motifs from individual CLIP datasets usually perform an enrichment analysis on a set of foreground and background sequences extracted from the CLIP data. For example, the mCross method derives foreground sequences from crosslinking peaks and background sequences from flanking regions upstream (−550…-450nt) and downstream (450…550nt) from the peak center [14]. Similarly, the ENCODE study derived foreground sequences from eCLIP peaks and background sequences from random areas within the same gene matched for length and transcript region [3, 15].

Analysis of crosslink-associated features, as compared to full sequencing reads, increases the accuracy of tools that identify enriched motifs [14, 16], because crosslink sites tend to coincide or be in close proximity of the RNA sequences that are recognized by the RBP. Nevertheless, motifs enriched at crosslink sites can also reflect technical biases of CLIP experiments, such as the sequence and structural preferences of UV-crosslinking [6], as well as the preferences of enzymes used for RNA fragmentation, ligation and reverse transcription [17], and sequence variations between genomic regions bound by different RBPs. The impact of these potential biases on motifs identified from CLIP data is poorly understood due to the lack of systematic studies [6].

Here we developed Positionally-Enriched k-mer Analysis (PEKA), a computational method that enables detailed examination of RBP’s binding preferences by visualising enriched motif groups for various RNA regions, which can be performed with or without considering the repetitive elements. To account for crosslinking biases and other technical features that affect the starts of reads in cDNA truncation-based methods (iCLIP and related) and transitions in PAR-CLIP [6], PEKA uses low-count crosslinks from the analysed dataset as a background. We show that PEKA performs equally well as mCross k-mer analysis when benchmarked using orthogonal *in vitro* data, while also allowing more flexible analysis tailored to the studied RBP. Here, we employ PEKA to perform comparative analysis of motif enrichment across 223 publicly available eCLIP datasets across 150 RBPs in addition to 3 iCLIP and 9 PAR-CLIP datasets for the same or closely related RBPs. We integrate it with orthogonal *in vitro* data to examine technical biases of eCLIP and to show that the extent of motif enrichment and inter-eCLIP motif specificity are useful signatures of data quality, and are linked to the domain composition, crosslinking efficiencies and regional binding preferences of the studied RBPs. To demonstrate that PEKA is an efficient tool for analysis of individual CLIP datasets, we did not perform any inter-RBP normalisation [14] or external background analysis [3].

By being able to derive reliable enriched motifs from analysis of individual CLIP datasets, PEKA will be useful to those performing CLIP experiments of any scale, and in any cell type or species. The source code for PEKA software is available at github [18], and the software is integrated into the iMAPS webserver [19], where it can be used for interactive and password-protected analysis of uploaded CLIP data. Moreover, we provide interactive analyses of eCLIP data in a variety of formats via a web-interface [20], including protein-centric view of motif enrichments, motif profiles around crosslinks of specific RBPs, and motif-centric analysis of comparative motif distributions around crosslinks of multiple RBPs. Such user-friendly discovery and exploratory visualisation of RNA binding motifs is a crucial step towards identifying the functionally-relevant binding sites from CLIP data as a basis for further studies.

## Results

### Positionally Enriched k-mer Analysis (PEKA) Overview

PEKA includes several features that are designed to examine and minimise the impact of technical biases of CLIP, and thereby to obtain enriched motifs that mediate RBP binding specificity. PEKA can perform motif discovery either across the full transcriptome or within defined transcriptomic regions, with the provided options including introns, 3 ′UTR, remaining exonic regions of protein-coding genes (coding sequence (CDS) combined with 5 ′UTR), non-coding RNAs (ncRNAs) and the rest of non-annotated intergenic regions (Figure 1a). PEKA also provides an option to include or exclude repetitive regions in the analysis.

**Figure 1.**
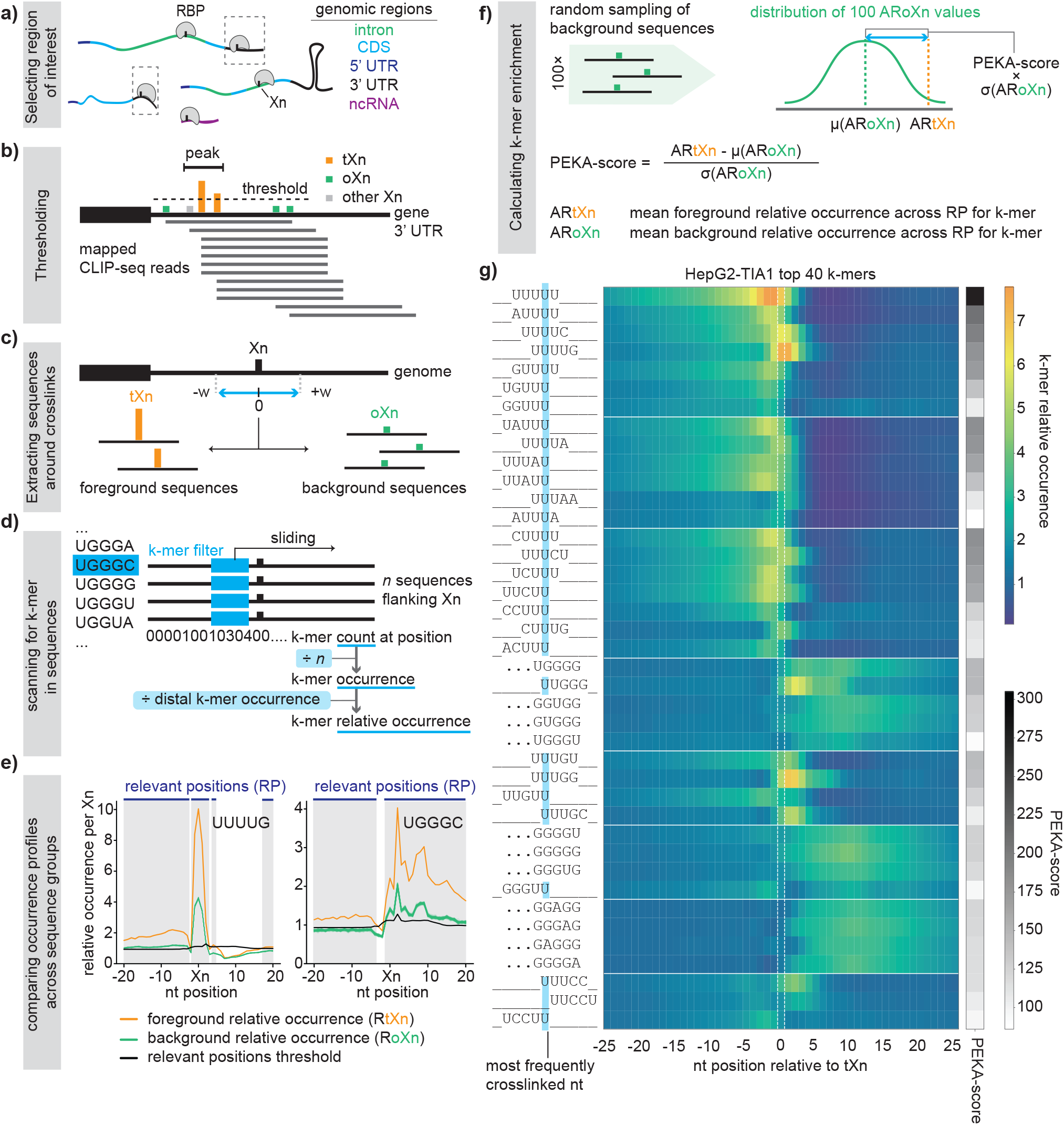
Schematic representation of the positionally enriched k-mer analysis (PEKA). **a)** Most eCLIP datasets have sufficient coverage on multiple transcriptomic regions to enable each to be investigated separately for motif enrichment by PEKA. **b)** Two bed files with the location of putative crosslink sites (Xn) and peaks are provided as inputs, so that PEKA can separate the Xn into the thresholded (tXn) and reference crosslink sites (oXn). Crosslinks that are not classified as tXn or oXn, are marked as other Xn. tXn lie within peaks and have a cDNA count equal or above a regional threshold, while oXn lie outside of peaks. **c)** Foreground and background sequences flanking tXn and oXn, respectively, are retrieved. The width (w) of the flanking region can be specified by the user. **d)** Sequences are scanned to record whether or not a k-mer is present at a particular position. For each group of sequences, k-mer occurrence around Xn is calculated and then converted to relative occurrence by dividing by mean k-mer occurrence across the distal region located 100 nt upstream and downstream from tXn (distal k-mer occurrence). **e)** Positions around Xn where relative occurrence at tXn passes the ‘position threshold value’ are considered as ‘relevant positions’ for enrichment analysis of each k-mer. **f)** PEKA-score is calculated for each k-mer by comparing the ‘mean k-mer relative occurrence’ across the relevant positions around tXn vs randomly sampled oXn, and used to rank the k-mers. **g)** Heatmaps showing relative occurrences (RtXn) and PEKA-scores for top 40 k-mers identified by PEKA for TIA1 eCLIP in HepG2 cell line. k-mers are clustered based on their sequences, their sequence on the left is aligned to the position of their maximal RtXn, and the most frequently crosslinked nucleotide is highlighted in blue. When the maximal occurrence position is located further from the crosslink site, three dots are shown.

Importantly, PEKA implements an approach of background normalisation that aims to minimise the technical biases at crosslink sites that are assigned by the starts of sequencing reads, which can include nucleotide preferences of UV crosslinking or sequence biases of cDNA ligation. To achieve this, PEKA extracts the background sequences that are centred on low-scoring crosslink sites (out-of-peak crosslinks, oXn), which are located outside the crosslinking peaks (i.e., areas with high crosslink density). The peak file is provided by the user, and peaks can be identified by any peak calling tool. For the purpose of current study, peaks have been identified with the Paraclu peak caller [21], largely due to its high speed and good performance in a previous iCLIP study [22]. Conversely, the foreground sequences are centered on high-scoring crosslink sites located inside the crosslinking peaks (thresholded crosslinks, tXn). The first step of PEKA is thus to split crosslink sites into thresholded (tXn) and out-of-peak (oXn) sites based on presence or absence of overlap with a peak file of choice, and on cDNA count at each crosslink with regards to a threshold that is defined in a region-specific manner (Figure 1b, and see Methods for more detail). Notably, tXn and oXn sites are expected to be affected by the same technical biases since they are part of the same cDNA library.

PEKA derives background and foreground sequences from user-defined genomic windows centred on tXn and oXn, respectively (Figure 1c). In the current study, these proximal windows were 41nt long, centered on the crosslink sites (i.e., ±20nt from Xn). The size of the proximal region can be adjusted to look for motifs closer or further away from the crosslink site. k-mer counts at each position across extracted sequences are transformed into k-mer occurrences by dividing them by the number of evaluated sequences. These are further transformed into ‘relative k-mer occurrences’ by dividing k-mer occurrence at each position within the proximal window with the average k-mer occurrence in a distal region (in this study defined as ±100- 150nt from tXn) (Figure 1d). A PEKA-score is then calculated by assessing the differential motif enrichment in the foreground relative to the background (Figure 1e,f). PEKA-score is a derivative of standard score and measures the number of standard deviations separating the estimated motif occurrence around tXn (ARtXn) from the mean estimated occurrence around oXn (μ(ARoXn)) (Figure 1f, and see Methods for details). As such, PEKA-score conveys the extent of k-mer enrichment relative to the internal background of the studied dataset. The motifs are then ranked based on PEKA-score in a descending order.

The top n k-mers are then selected to visualise their positioning around tXn with occurrence profiles. By default, PEKA separates the top 20 k-mers into up to 5 clusters based on their similarity in sequence and occurrence profiles, and then visualizes the profiles of k-mers within each cluster on the same graph (Suppl. figure 1, Additional file 1). For heatmap visualisations, the top 40 k-mers are clustered based on their sequence, and their relative occurrence profiles are shown (Figure 1g). The use of relative occurrence, which normalises the raw k-mer occurrences with distal region (Figure 1d), improves the capacity to compare the enriched positions of different k-mers, as otherwise the regional genomic differences of k-mers would decrease the visibility of the less-abundant k-mers.

### PEKA enables combined analysis of motif enrichment and positioning

To get preliminary insights into the distribution of the enriched k-mers around crosslink sites (Suppl Figure 1), we first analysed the occurrence profiles of 5-mers for the well-studied protein TIA1. The 5-mer profiles for TIA1 eCLIP in the HepG2 cell line show the most enriched motifs to be U-rich (Figure 1g), in agreement with the known sequence specificity of TIA proteins [23]. In particular, the top 12 k-mers are all U-rich, followed by ‘UGGGG’ and several other G-rich k-mers. Interestingly, U-rich motifs are enriched directly at crosslink sites, whereas G-rich motifs are depleted at crosslink sites and enriched primarily downstream of the crosslink sites. These non-overlapping positional patterns of the two groups of motifs indicate either that they represent different modes of binding by TIA1, or that the G-rich motifs are derived from other co-purified RBPs that bind to the G-rich motifs. This case-study demonstrates first that k-mers with similar sequences tend to have similar positional profiles around crosslink sites; and second, the ability of PEKA to report multiple motif types with distinct profiles, which can yield insights into data specificity or multiple binding modes of RBPs. Given the useful insights derived from this visualisation, we provide the heatmaps of the top 40 k-mers determined by PEKA for all 223 ENCODE eCLIP datasets on the web-interface [20].

### Benchmarking of PEKA against mCross and *in vitro* data

A major challenge in CLIP data analysis is the identification of motifs that contribute to the RNA-binding specificity of the isolated RBP, as opposed to motifs enriched for other confounding reasons, such as technical biases of CLIP, associated features of genomic regions and repetitive elements bound by the protein, or motifs bound by other RBPs that may be co-purified if IP stringency was insufficient [6]. To evaluate the specificity of enriched motifs, these can be cross-compared with the motifs obtained by *in vitro* methods, in particular the high-throughput methods RNA-Bind-n-Seq (RBNS) [24] and RNA-compete (RNAC) [25]. In both RNAC and RBNS, a specific RBP or an assembly of its RNA-binding domains is mixed with a random pool of RNAs, followed by the isolation and sequencing of bound RNA molecules. These *in vitro* methods are expected to have different biases than CLIP data, and their limitations arise mainly from the use of short RNA fragments and the absence of full-length proteins and their cofactors. Thus, data produced with these methods is well suited to examine the biological specificity of motifs derived from CLIP data.

To examine the capacity of PEKA to recognise the top-ranking motifs obtained from *in vitro* data, we compared the top 20 k-mers recovered from *in vitro* RBNS/RNAC datasets with the motifs identified by PEKA in the corresponding eCLIP data (Suppl. Figure 2a, Additional file 1). We also compared the performance of PEKA with that of a previously published mCross method on the same data [14]. In total, 41 eCLIP datasets (Additional file 2) were compared for 28 distinct proteins as they contain both mCross and *in vitro* data. As RBNS provides 5-mer scores, PEKA was also run for 5-mers, and to ensure a valid comparison, the mCross and RNAC z-scores were converted from 7-mer to 5-mer scores prior to ranking the motifs. Ranking k-mers by normalized mCross 5-mer z-scores resulted in k-mer logos (Suppl. Figure 2b, Additional file 1) that were very similar to the mCross sequence logos downloaded from the mCrossBase [14, 26] (Suppl. Figure 2c, Additional file 1), confirming that the relevant k-mers were retained during this conversion. Importantly, PEKA overall showed a similar performance to mCross in recovering the top 20 k-mers from *in vitro* data (Suppl. Figure 2a, Additional file 1).

To compare PEKA and mCross in more detail, we returned to the G-rich motifs observed for TIA1 eCLIP in HepG2 cell line (Figure 1g). Because these motifs are unexpected, and have not previously been reported to be bound by TIA1, they could be considered a potential artifact of PEKA. Therefore, we performed a direct comparison of the top 20 5-mers obtained for TIA1 eCLIP in HepG2 cell line, using either PEKA-score, raw mCross z-scores or normalized mCross z-scores, respectively. Based on our observation of the two k-mer groups in the data (U-rich and G-rich, Figure 1g), the top 5-mers were divided into two clusters based on their sequence and visualized as k-mer logos (Suppl. Figure 2b). Reassuringly, G-rich k-mers were also detected by the raw mCross z-scores to a similar extent as by the PEKA-scores. However, they were largely removed in the normalised mCross z-scores, which penalise the k-mers with high z-scores in most experiments [14]. Thus, detection of G-rich k-mers in TIA1 eCLIP by PEKA is valid, and such k-mers could be enriched in multiple eCLIP experiments.

### Insights from variable motif enrichments across CLIP variants

To learn more about the motif preferences of different CLIP protocols, we performed a more detailed comparison of PEKA with the results obtained by *in vitro* method RBNS [3], which are available along with eCLIP data for 21 RBPs, iCLIP data for 3 of these RBPs, and PAR-CLIP data for 8 RBPs (Additional file 2,3). PEKA analysis was performed in the ‘protein-coding gene’ region, which combines intron, CDS and the untranslated regions (UTRs). Repeat sequences were filtered out in motif detection. Visualisation of k-mer ranking across groups, generated by sequence-based clustering, forms a unique binding signature for each dataset (Figure 2). We observed informative variations in the ranking of top k-mers between data produced by these three CLIP methods.

**Figure 2.**
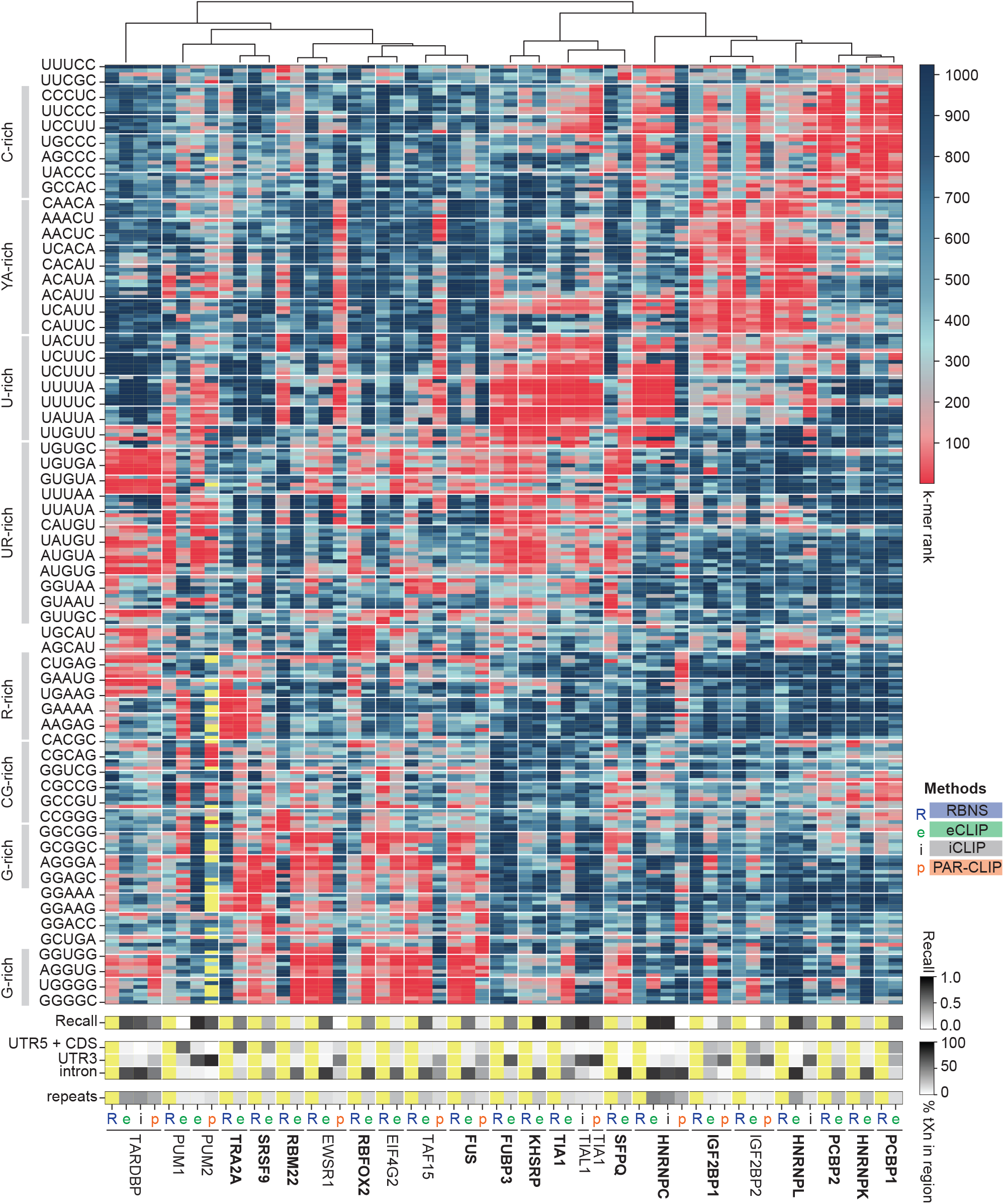
Comparison of CLIP methods against RBNS. Heatmap shows the rank order of k-mer PEKA-scores for each CLIP dataset, and of RBNS z-scores. The scale on the top right shows these values spanning from 1 to 1024. k-mers selected for this heatmap (n=245) ranked among the top 10 in eCLIP, PAR-CLIP, iCLIP or RBNS for any of the RBPs shown. k-mers were clustered based on their sequence and ordered using hierarchical clustering of median dataset ranks within clusters. On the left, prominent k-mer sequence features are highlighted across clusters. Below, the recall heatmap shows how much the top motifs in the dataset correspond with the RBNS data. The third heatmap shows the percentage of tXn derived from different transcript regions, and the fourth heatmap shows the percentage of tXn found in repeat sequences. eCLIP experiments in bold are in HepG2, others in K562 cell line. In case two eCLIP datasets for the same RBP were available, the one with higher recall is shown.

In the case of TARDBP, FUBP3, KHSRP, TIA1, HNRNPC, HNRNPL, PCBP1 and PCBP2, eCLIP as well as most of the available iCLIP and PAR-CLIP experiments show high agreement with RBNS (i.e., recall, Figure 2). eCLIP has a slightly lower agreement with RBNS than iCLIP for TIA1, which is mainly due to the presence of G-rich motifs in eCLIP but not in TIA1 iCLIP, PAR-CLIP or RBNS. The G-rich motifs are also enriched in RBFOX2 and IGF2BP1/2 eCLIP, but not in RBNS, or IGF2BP1/2 PAR-CLIP. In the case of IGF2BP1/2, additional divergence can also be attributed to the enrichment of C-rich motifs in eCLIP, which RBNS and PAR-CLIP do not exhibit. In addition to IGF2BP1/2, eCLIP experiments for PUM1, RBM22, and EIF4G2 also showed poor agreement with RBNS data. In the case of PUM1 and IGF2BP1/2, the data from PAR-CLIP are in much better agreement with RBNS than eCLIP. Considering the known similarity of motif specificity of PUM1 and PUM2 [27], we compared the PUM2 eCLIP experiment with the *in vitro* data of PUM1, which showed high agreement, as reported previously [3]. This analysis demonstrates the large differences in the reliability of eCLIP datasets, and the value of CLIP meta-analyses to identify the datasets that are likely to be the most reliable for further studies.

We found that when CLIP datasets differ in enriched motifs, they tend to also differ in the regional distribution of crosslink sites (Figure 2). Specifically, compared to eCLIP and iCLIP, PAR-CLIP has a generally higher proportion of tXn in the 3 ′UTR relative to introns, and a lower coverage of repetitive elements. tXn distribution also varied between eCLIPs of homologous proteins PUM1 and PUM2, where we found PUM 1 to have a lower proportion of tXn in the 3 ′UTR and a higher proportion of intronic tXn. Thus, a combined analysis of several CLIP features, such as motif enrichment and regional binding, may be particularly valuable for data quality assessment, and for understanding the potential generic biases of each CLIP variant.

### Benefit of peak filtering by external background is limited

When analysing eCLIP data it is important to consider that the ENCODE consortium provides narrowPeaks generated by filtering the data to retain only those peaks with a significant enrichment of reads over the a size-matched input (SMInput) control [5]. Such filtering is expected to decrease the extent of extrinsic background in data [3,6,28]. In contrast, PEKA was developed for general use on any type of public CLIP data, and therefore it models motif enrichment relative to the intrinsic background. To understand the importance of using SMInput, we additionally ran PEKA by using only the crosslink sites located within the narrowPeaks as the foreground. Strikingly, we observed no overall improvement in motif specificity when comparing PEKA motifs identified when using raw eCLIP data as input to the standard Paraclu peak calling approach to identify tXn, or when using only tXn from narrowPeaks (Figure 3a). A major increase in agreement with *in vitro* data when using narrowPeaks was found only for a few specific RBPs, namely IGF2BP1 and SRSF1, but a major decreased agreement was seen for many other RBPs, such as hnRNPC, PTBP1, RBM5 and hnRNPA1.

**Figure 3.**
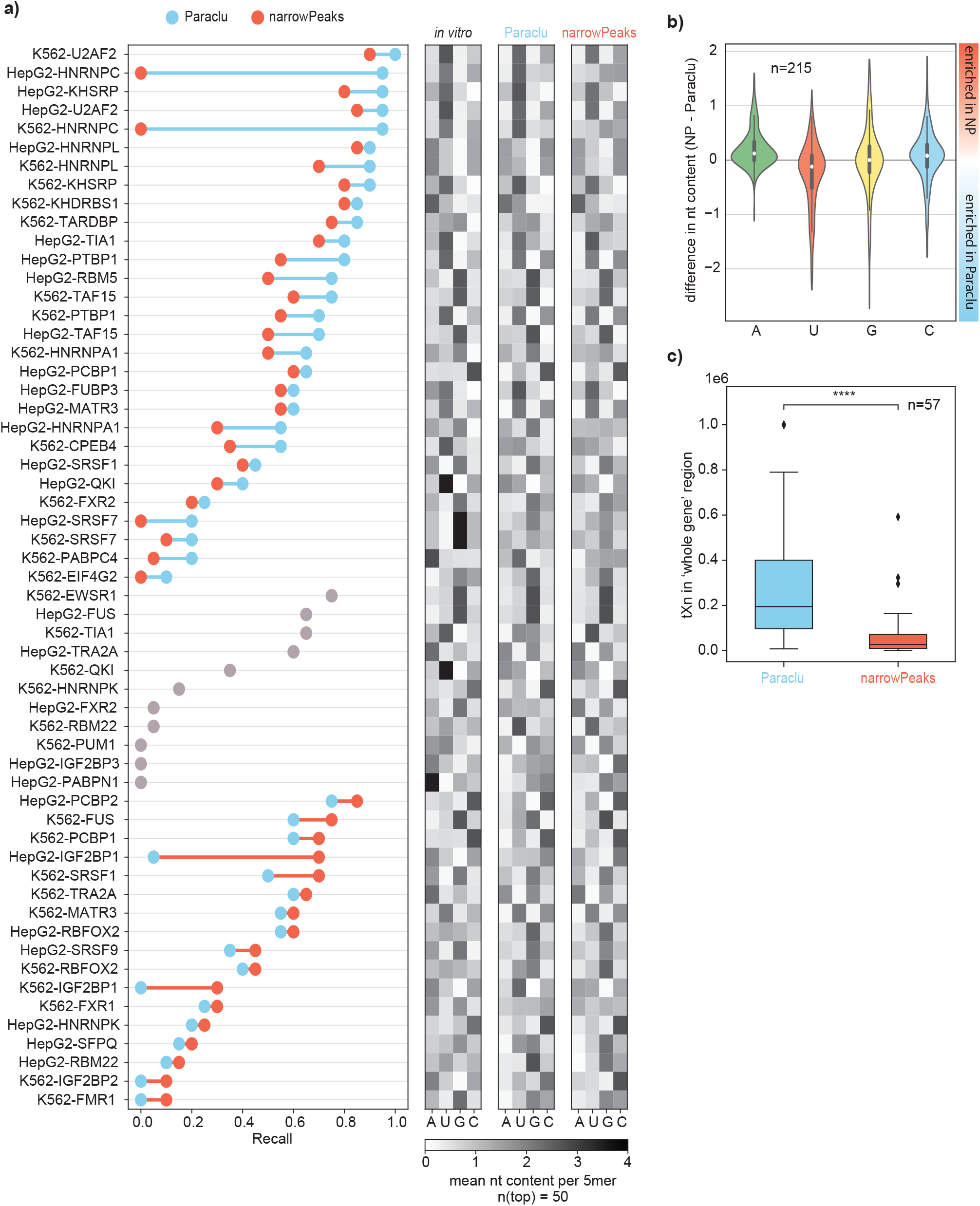
Influence of size-matched input controls on motif discovery. **a)** The graph shows recall for 57 eCLIP datasets for which orthogonal RBNS or RNAcompete data was available. The datasets were processed with PEKA, using either Paraclu peaks or eCLIP narrowPeaks from merged replicates, downloaded from the ENCODE consortium [4, 5]. The datasets are sorted into three groups: 1) datasets where higher recall was achieved with Paraclu peaks, 2) datasets with no change in recall between groups and 3) datasets where higher recall was achieved with narrowPeaks. Within each group datasets are ordered by decreasing recall values. Heatmaps on the right show mean nucleotide composition across the top 50 k-mers as ranked by *in vitro* method, by PEKA using Paraclu or PEKA using narrowPeaks. **b)** Violin plots show differences in mean nt composition of top 50 k-mers for 215 eCLIP datasets, which had sufficient tXn coverage in narrowPeaks for PEKA analysis. **c)** Number of tXn detected by PEKA in the ‘protein-coding gene’ region when using Paraclu vs narrowPeaks (paired t-test, p < 0.0001) for 57 eCLIP datasets shown in panel a).

We find that the approach used to derive narrowPeaks generally results in a decrease in the proportion of crosslinks that identify binding to uridine-rich motifs and an increase the proportion of crosslinks that identify motifs containing adenosine and/or citidine (Figure 3b). As a result, the use of narrowPeaks tends to improve motif specificity for RBPs that do not show binding to U-rich motifs as defined by *in vitro* data, but it often decreases specificity for proteins that bind to U-rich motifs (Figure 3a). For example, IGF2BP1 primarily binds to AC-rich motifs, which are enhanced with narrowPeaks. To better understand this phenomenon, we applied PEKA separately to the eCLIP and SMInput dataset for hnRNPC, where the largest decrease in specificity is observed with narrowPeaks. Interestingly, we find that U-rich motifs are most strongly enriched both in hnRNPC eCLIP and SMInput, with a very similar profile around crosslink sites in both cases, indicating that crosslink sites of hnRNPC likely represent part of the SMInput signal (Suppl. Figure 3a,b, Additional file 1).

Since we observed that G-rich motifs are most frequently enriched in eCLIP datasets, we investigated how narrowPeaks impact the enrichment of G-rich motifs in TIA1 eCLIP (Figure 1g). For this we compared their ranking by PEKA in combination withParaclu peaks or narrowPeaks, respectively. We found that only 3 k-mers with ≥ 3 guanosines are among the top 40 k-mers when narrowPeaks is used, but 13 are found when Paraclu is used. This argues that the use of narrowPeaks decreases the detection of G-rich motifs. Interestingly, however, recall of TIA1 HepG2 eCLIP compared with *in vitro* data was still slightly higher for Paraclu compared with narrowPeaks when used together with PEKA (Figure 3a), indicating that the decreased enrichment of G-rich motifs did not result in higher enrichment of relevant k-mers.

We also found that the median number of tXn that overlap with narrowPeaks is around 7-fold lower compared to the Paraclu peaks used with PEKA (Figure 3c). Thus, the approach of SMInput filtering, which was used to identify narrowPeaks, greatly reduces the sensitivity of data without generally improving the specificity of motifs identified by PEKA. Moreover, crosslink sites located in narrowPeaks are biased away from U-rich motifs, even for proteins known to bind such motifs, and toward A-rich motifs, which could lead to systemic biases. Taken together, this analysis demonstrates that the value of SMInput-based correction depends on the specificity of the studied RBP, which might reflect contributions of certain RBPs to the SMInput data. As a result, for proteins such as hnRNPC SMInput likely represent a mixture of foreground and background signal, leading to depletion of the foreground signal from narrowPeaks. Such loss of relevant foreground signal can be particularly detrimental when used together with PEKA, as lost foreground signal will be modeled as background, causing a decreased enrichment of relevant motifs. Taken together, this analysis demonstrates that it is appropriate for PEKA to perform motif analysis by relying on the intrinsic background modeled from each CLIP dataset individually, instead of using SMInput filtering.

### Insights into technical biases of CLIP

To analyse potential systematic differences between various methods that identify enriched motifs, we compared the motif enrichment between PEKA, mCross and RBNS across a subset of 24 eCLIP datasets (representing 17 distinct RBPs, Additional file 2) that had both RBNS and mCross data available. To illustrate how PEKA controls for technical biases, we compared the motif ranking of PEKA and mCross with a simple ‘local’ approach that only examines the local motif occurrence around tXn, without normalizing by the intrinsic background (i.e., by the relative occurrence around oXn). This ‘local approach’ ranked the k-mers by their maximal value of relative occurrence in the −25…25 window around tXn.

We first identified groups of k-mers that were differentially ranked in each approach of eCLIP analysis relative to RBNS (confidence interval > 95%) (Figure 4a, see methods for details). In this way we wished to gain insights into the reasons why certain motifs are more or less recoverable from eCLIP than *in vitro* data. Interestingly, the ‘local’ approach, which does not correct for technical biases, produced most motifs that were differentially enriched in eCLIP vs RBNS (30), and PEKA produced the least (12). Notably, only two motifs were differentially enriched in eCLIP by all three approaches (CGUCU and UUCGU). Conversely, 8 motifs were more enriched in RBNS compared to all three approaches to eCLIP analysis. These findings highlight the importance, as well as the robustness of the intrinsic background correction performed by PEKA.

**Figure 4.**
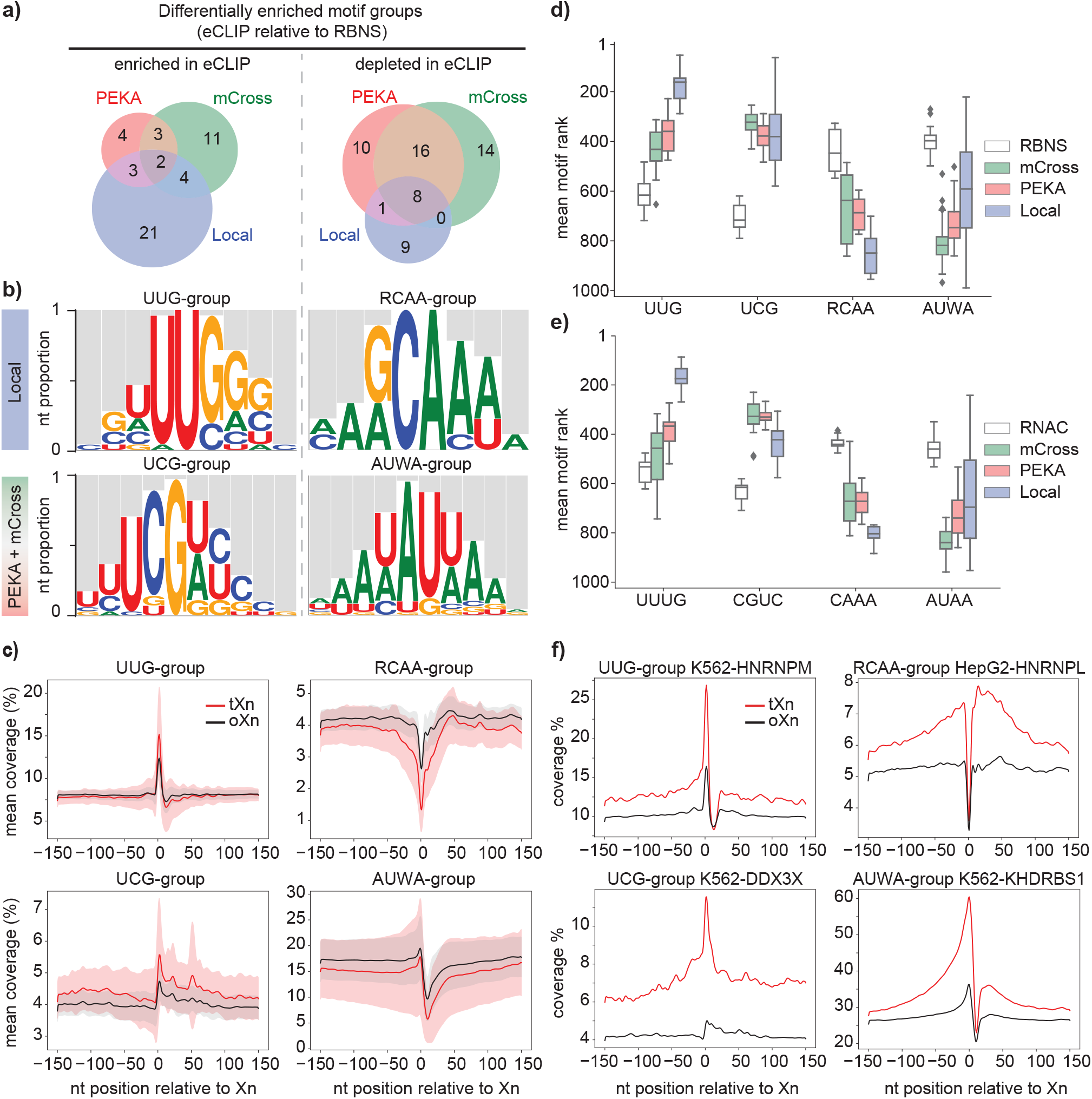
Differential enrichment of motif groups in eCLIP relative to in vitro methods. **a)** Venn-diagrams show the overlap across groups of 5-mers with significant difference in ranking between RBNS and eCLIP, analysed with PEKA, mCross and ‘local’ approach. **b)** k-mer logos for groups of differentially ranked 5-mers (see Methods) between RBNS and eCLIP, analysed with either by PEKA or mCross or by ‘local’ approach. Nucleotides shown in k-mer logos are scaled proportionally to their frequency at each position in the alignment. **c)** Distribution of mean motif-group coverage around tXn (red) or oXn (black) within the ‘protein-coding gene’ region, across 24 eCLIPs with orthogonal RBNS data. Area around the curve shows the standard deviation. **d, e)** Boxplot showing distribution of mean motif ranks across RBPs between RBNS (d) or RNAC (e) and methods used to analyze eCLIP for the four motif-groups. **f)** Distribution of mean motif-group coverage around tXn (red) or oXn (black) within the ‘protein-coding gene’ region for RBPs that are known to bind these motif groups.

Both PEKA and mCross use a strategy to control the technical biases of CLIP and interestingly, the differential motifs of both methods share more similar features, such as UC- and CG-containing motifs, with each other than with the ‘local approach’ (Suppl. Figure 4, Additional file 1). Thus, to understand the extent to which the two methods correct the technical biases, we combined the differential motifs from both methods for further analyses, and compared these motifs to those that were differentially enriched or depleted in the ‘local’ approach, but not in PEKA or mCross. We started by analyzing a group of k-mers (n=21) that were differentially enriched only in the ‘local approach’ for the 24 eCLIP datasets, as compared to RBNS. As expected from the known crosslinking bias of U and G, these k-mers are highly U/G-rich (UUG-group, Figure 4b), and their peak of coverage directly aligns to the crosslinking sites (Figure 4c). Importantly, while these k-mers are highly enriched when using the ‘local’ approach, their median rank drops to below 300 in PEKA and mCross (Figure 4d), demonstrating that both methods are quite effective in diminishing this technical bias of CLIP. To corroborate these insights, we repeated the same differential motif analysis by comparing k-mer ranks between RNAcompete and eCLIP on 27 datasets, representing 19 distinct RBPs (Additional file 2). This yielded motif groups with similar sequence features (Figure 4e), indicating that the differential motifs stem from general differences between the eCLIP method and the *in vitro* methods.

The reason for the capacity of PEKA to diminish the bias of U/G-rich kmers can be seen by analysing their occurrence at tXn vs oXn across the 24 datasets (Figure 4b). Clearly, the peak of U/G-rich k-mers is high at both types of crosslink sites, and therefore normalisation by occurrence at oXn decreases PEKA-score (see Figure 1f) and consequently increases the ranking of these motifs. Crucially, however, these motifs remain top-ranking for RBPs that truly bind them with high affinity, as is exemplified by hnRNP M [29], where the enrichment around tXn is much higher than at oXn (Figure 4f). Thus, PEKA is able to distinguish motifs that are enriched at all crosslink sites due to crosslinking biases from those that mediate high-affinity RBP binding, as they are enriched at crosslink sites with high cDNA counts (i.e., at tXn).

We next analyzed the group of k-mers that was most differentially depleted in the ‘local approach’ for the 24 eCLIP datasets, as compared to RBNS. These k-mers are depleted of Us and rarely contain a G, and align with the RCAA motif (Figure 4b). They are strongly depleted at the crosslink sites (as compared to the surrounding region), which most likely reflects a negative crosslinking bias (Figure 4c). Low crosslinking of these motifs would be consistent with the prevalence of A and C, nucleotides that don’t crosslink well. However, the median rank of these motifs increases in PEKA and mCross as compared to the ‘local approach’ (Figure 4d), and thus both methods partly diminish this potential technical bias of CLIP. Importantly, PEKA can identify these motifs as top-ranking for RBPs that bind them with high affinity, as is exemplified by hnRNP L [30], where the enrichment around tXn is much higher than at oXn (Figure 4f). Thus, in spite of the low crosslinking efficiency of A/C-rich motifs, CLIP can identify the binding sites containing such motifs, in which case the motifs are enriched in close proximity of crosslink sites that have high cDNA counts (i.e., at tXn).

Next, for the 24 eCLIP datasets we investigated the group of k-mers that was most differentially enriched in PEKA/mCross as compared to RBNS. These k-mers align with the UCG motif (UCG-group, Figure 4b). They are enriched to a similar extent (compared to oXn) up to 150nt around tXn (Figure 4c). Interestingly, the DEAD-box helicase DDX3X is among the proteins that rank most highly for these k-mers, which are broadly enriched around tXn crosslink sites (Figure 4f). DDX3X contains IDRs with strong condensation propensity and is a major regulator of cellular RNA condensates [31]. A study also reported that DDX3X binds in vicinity to a motif composed in large part of CG and CGU subsequences, similar to the k-mers found in UCG-group [32].

Finally, we studied the group of k-mers that was most differentially depleted in the PEKA/mCross for the 24 eCLIP datasets, as compared to RBNS. These k-mers align with the AUWA motif (Figure 4b). They are depleted to a similar extent (compared to oXn) up to 150nt around tXn (Figure 4c). PEKA nevertheless identifies these k-mers as top-ranking for RBPs that bind them with high affinity, as is exemplified by Khdrbs1 [33], where the enrichment around tXn is much higher than at oXn (Figure 4f).

We also noted that the enrichment of UCG-group, and depletion of RCAA-group and AUWA-group motifs around tXn (as compared to oXn) extends far from the region containing the CLIP reads (just downstream of crosslink sites) (Figure 4c). Thus, it is unlikely just a result of technical biases of CLIP, such as sequence biases of crosslinking, RNase and ligation reactions. Instead, it is plausible that these broader patterns reflect the preferences of the studied RBPs for motifs distributed across broad RNA regions. This would be consistent with the recent report that condensation-dependent assembly of TDP-43 occurs on RNA regions containing broad clusters of binding motifs [22], and that regions with clustered motifs have increased probability of Dazl binding [34].

### Many eCLIP datasets share similar enriched motifs

To understand how specific are the motifs enriched in each eCLIP dataset, we performed a systematic cross-motif and cross-RBP comparison of the whole set of eCLIP datasets available from ENCODE [4, 5]. We clustered the data based on the regional crosslinking profiles within protein-coding genes (CDS/UTR/intron) and by the ranks of 5-mers (see Methods), which identified groups of eCLIPs with similar motif enrichment signatures (Figure 5). The first visually apparent feature of this analysis is that regional crosslinking preferences are accompanied by trends towards certain motif preferences. For instance, if crosslinks are primarily in 5 ′UTR and CDS, the largest cluster of data shows enrichment in purine-rich motifs. In case of datasets with primary crosslinking in introns, motif enrichment is dominated by two clusters, a very large cluster of datasets dominated by G-rich motifs, and a smaller cluster dominated by U-rich motifs. Datasets that crosslink primarily to 3 ′UTRs also show enrichment of U-rich or UA-rich motifs (see the blue and yellow/red cluster, respectively). These motifs likely include binders of the AU-rich elements (AREs) that are common regulators of RNA stability in 3 ′UTRs [35]. Moreover, both CDS and intronic datasets are often enriched in C/G-rich motifs. This analysis demonstrates that the common motif preferences in eCLIP data are closely linked to the regional crosslinking profiles within the protein-coding genes.

**Figure 5.**
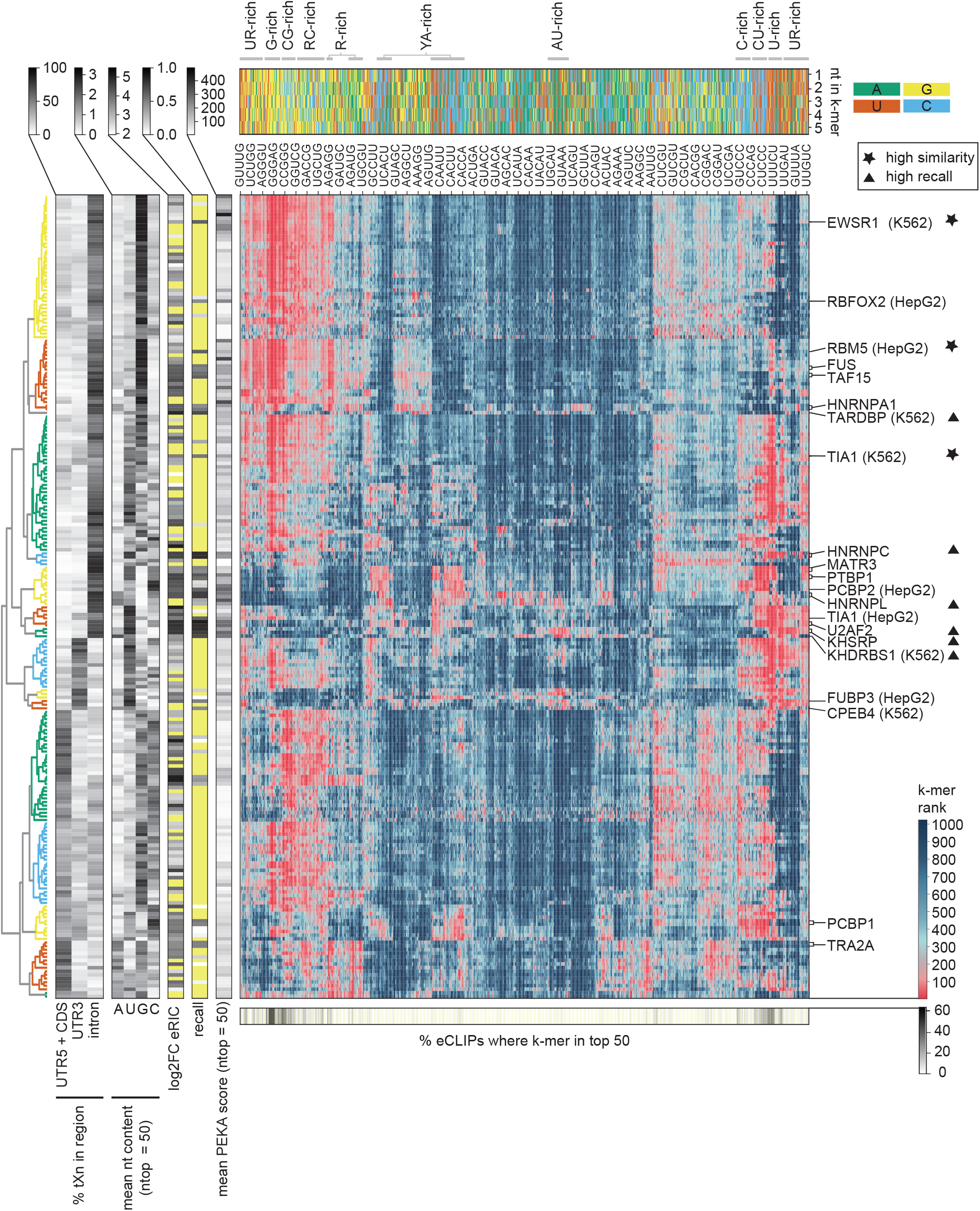
Heatmap of all 5-mers for all available eCLIP datasets. eCLIP datasets are hierarchically clustered by their rank order of 5-mers and regional distribution of high-confidence crosslink sites (tXn) to visualize binding preferences across groups of proteins. 15 primary clusters were identified, which are colour coded in the dendrogram on the left. Additional heatmaps on the left represent (from left to right) 1) regional distribution of thresholded crosslink sites, 2) mean nucleotide composition across top 50 identified k-mers for each dataset, 3) enrichment of the protein in the RNA interactome capture (eRIC [36]), 4) the overlap with orthogonal *in vitro* methods (i.e. recall) and 5) mean PEKA-score across top 50 identified k-mers. RBPs marked on the main heatmap have a recall value > 0.5. A triangle next to the RBP name represents high recall (> 0.8) and a star represents high similarity score (> 0.3). Additional heatmap above the main heatmap represents the nucleotide sequence of each 5-mer and a grayscale heatmap below the main heatmap shows a percentage of eCLIPs, where k-mer ranked among the top 50. Every 20-th 5-mer is labeled on the main heatmap. Yellow fields in grayscale heatmaps indicate missing values.

As expected, eCLIP datasets within the largest clusters share highly similar motif preferences and regional profiles. Interestingly, we noticed that RBPs falling within these large clusters generally lack orthogonal *in vitro* binding data (as seen by the absence of recall), are often poorly detected in the mRNA interactome proteomics (enhanced RNA-interactome capture i.e., eRIC) [36], and their top 50 ranked k-mers are often G-, U- or GC-rich. It has been reported previously that such k-mers tend to be overrepresented in eCLIP compared to RBNS [3], but the scale of their presence across eCLIP data was not yet examined. Strikingly, several G-rich k-mers were enriched among the top 50 k-mers in more than 50% of all eCLIP datasets.

### Specificity of eCLIP datasets relates to RBP domain types and compositional biases

To further understand how data similarity relates to the various features of RBPs, we divided the eCLIP datasets according to whether or not they had available *in vitro* data, and then clustered each group based either on a combination of inter-data similarity (similarity score, see Methods) and recall, or just on inter-data similarity where no *in vitro* data was available, obtaining 7 groups of data (Figure 6a and see Methods for details). Most apparent differences are seen between the group of proteins for which *in vitro* data are available (groups 1-4, i.e. ‘*in vitro* set’) and the group for which no data are available (groups 5-7, i.e. ‘eCLIP-only set’). The *in vitro* set tends to have high eRIC values and high mean PEKA-scores across the top 50 ranked k-mers, whereas the eCLIP-only set tends to have low eRIC values and low mean PEKA-scores, indicating that the proteins in the eCLIP-only set don’t crosslink well, and have low extent of motif enrichment, respectively. The great majority of proteins in the *in vitro* set contain a KH or RRM domain and a low-complexity sequence, which are the common characteristics of RBPs [37]. However, proteins in the eCLIP-only set rarely contain these features. Interestingly, the *in vitro* set contains only a small group 4 with high similarity score, while the eCLIP-only set contains a large group 7 with high similarity score, accounting for 40% of all eCLIP experiments. Taken together, it is clear that proteins lacking orthogonal *in vitro* data generally have different features from the rest, and their eCLIP data tends to have lower inter-data specificity (high similarity index) and motif enrichment (low mean PEKA-score, eCLIP-only set). This indicates that cross-validation of eCLIP with *in vitro* data cannot be extrapolated to warrant the specificity of eCLIP data for which there are no *in vitro* data, which must be be taken into account when performing meta-analyses on the whole set of eCLIP data.

**Figure 6.**
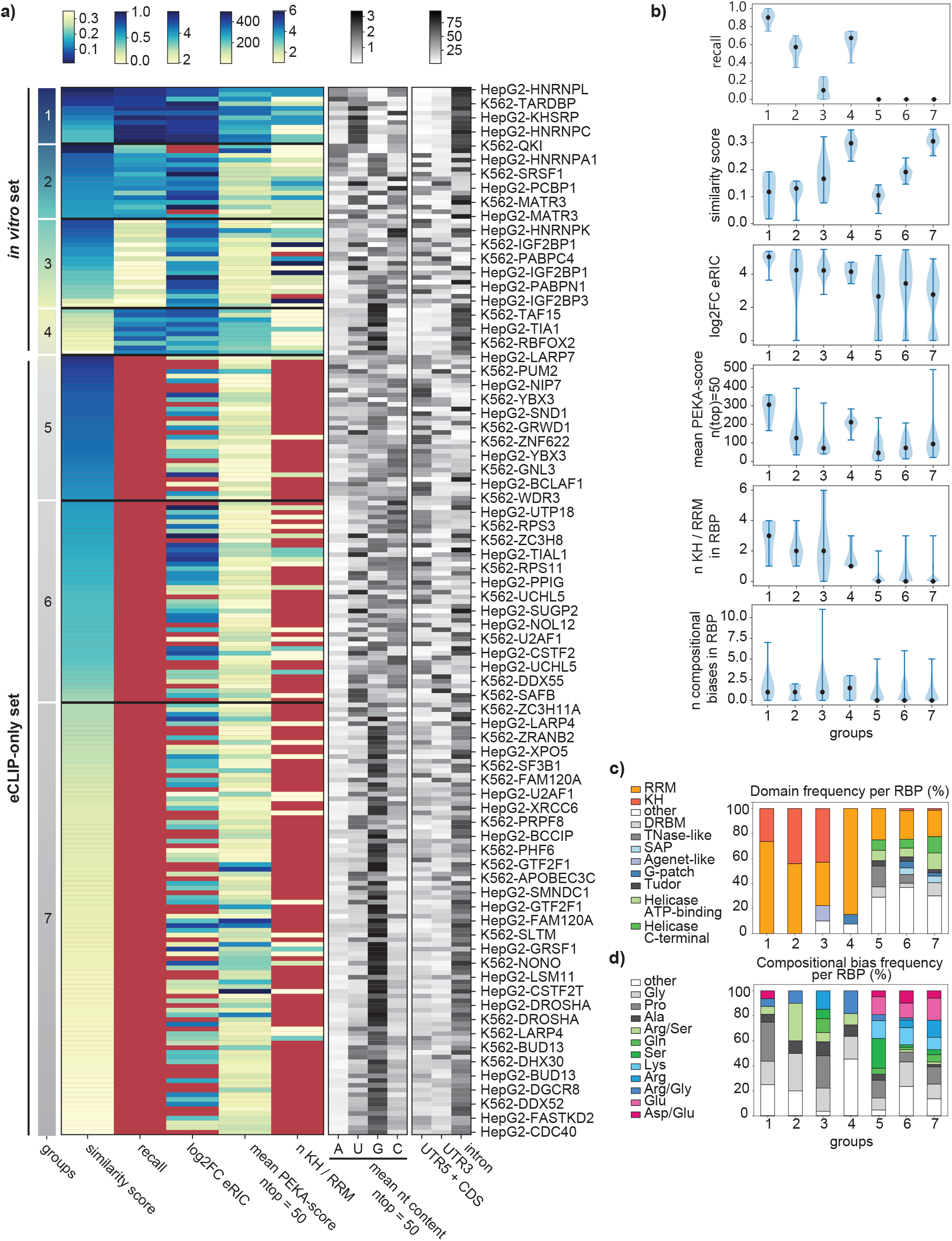
Expected specificity of eCLIP datasets with regards to various features. **a)** The left heatmap displays all eCLIP datasets, clustered based on their similarity scores and recall into 7 clusters. For each dataset the heatmap also shows its enrichment in the mRNA interactome proteomics (log2FC eRIC), the number of KH or RRM in RBP and the average PEKA-score across the top 50 ranked k-mers. Values shown on this heatmap are available in Additional file 7. Heatmaps on the right show the mean nucleotide content across the top 50 ranked 5-mers for each dataset and % of thresholded crosslinks derived from each transcript region. **b)** Violin plots showing (from top to bottom) 1) similarity score, 2) log2FC eRIC, 3) mean PEKA-score across top 50 k-mers, 4) the number of total domains in RBPs, 5) the number of KH or RRM domains for RBPs and 6) the total number of compositional biases for RBPs within each cluster. **c, d)** Stacked barplots showing the frequency of a particular domain (c) and compositional biases (d) per RBP within each cluster. Compositional biases are color-coded based on representative amino acid properties, with aliphatic amino acids in shades of grey, hydroxyl or amide functional groups containing amino acids in green, basic amino acids in blue and acidic amino acids in pink.

The most reliable eCLIP experiments are expected to be in group 1, which includes ∼5% of datasets with unique k-mer signatures and high agreement with corresponding RBNS or RNAC data, as indicated by their low similarity index and high recall, respectively. This group of RBPs generally ranked highest in the eRIC experiments, indicating that they crosslink efficiently with RNA. These RBPs contain a median of 3 RNA recognition motif (RRM) or K-homology (KH) domains (Figure 6b). Thus, the canonical RBPs that crosslink well and contain many RNA binding domains tend to yield specific and reliable eCLIP datasets. In addition to the high cross-validation, RBPs in group 1 have the highest mean PEKA-scores across the top 50 ranked k-mers, implying that the coverage of top k-mers around tXn is much higher than around oXn. In other words, binding affinity of these RBPs is strongly sequence-dependent, requiring the presence of one or more high-affinity binding motifs.

The least reliable eCLIP experiments are expected within group 7, containing ∼40% of eCLIP datasets with high inter-data similarity, which lack orthogonal *in vitro* data (Figure 6a). ∼36% of datasets in group 7 are undetected in eRIC experiments, which indicates that they crosslink poorly or do not crosslink at all to RNA. These proteins lack annotated features of RBPs, such as KH or RRM domains (Figure 6a,b). This increases the likelihood of the signal being dominated by the most common contaminants of eCLIP experiments, which are likely the abundant and well-crosslinking RBPs. Low-specificity datasets in group 7 predominantly enrich for G-rich motifs, with most crosslinking sites originating from introns (Figure 6a). We note that the same features are also prevalent in group 4, which contains eCLIP datasets that have reasonable agreement with *in vitro* data, indicating that many of these RBPs directly interact with the G-rich motifs. Nevertheless, these experiments have high inter-data similarity due to the fact that very similar motifs are enriched in group 4 and 7. However, RBPs in group 4 generally have much higher mean PEKA-scores than those in group 7, and thus even though both show enrichment of similar motifs, the extent of enrichment is stronger in group 4 (Figure 6b).

It is interesting to find many groups with strong regional binding preferences, even though regional preferences had no direct role in clustering the groups (Figure 6a). For example, groups 1, 4 and 7 all contain predominantly intronic binding. However, G-rich motifs dominate groups 4 and 7, whereas group 1 mainly shows enrichment of A-rich or U-rich motifs, likely due to its higher data specificity. It is notable that proteins in group 1 contain a median of 3 KH or RRM domains, whereas those in group 4 contain only a median of 1 domain, and don’t ever contain a KH domain. Moreover, group 5 commonly shows predominant binding in CDS and 5 ′UTR, which tends to be associated with a higher proportion of A-rich motifs. Since A-rich motifs are otherwise rare in eCLIP experiments, their enrichment contributes to the low similarity index of datasets in group 5. We propose two possible explanations why datasets with similar specificity tend to have similar regional binding. First, the signature might reflect similar contaminants; for example, RBPs that bind G-rich motifs in introns might be the common contaminants of datasets from group 7. Second, the link between RBP specificity and regional binding could reflect regional sequence biases.

In addition to the aforementioned differences in the content of RRM/KH domains, we found that our clusters of proteins also differed in other domains. The proteins in the *in vitro* set very rarely contain domains other than RRM/KH, whereas the proteins in the eCLIP-only set frequently contain helicase domains, TNase-like domain, Tudor domain and dsRNA-binding domains (DRBM) (Figure 6c). These domains are less sequence specific, which likely contributes to the generally low eRIC scores of these proteins, and the high similarity scores and low PEKA-scores of their eCLIP data.

As mentioned earlier, we found that proteins in the *in vitro* set tend to contain at least one LCR, which is not the case for RBPs in the eCLIP only set (Figure 6b). We also observed large differences in the types of compositional biases between the *in vitro* and eCLIP-only sets (Figure 6d). Proteins in the *in vitro* set contain a higher proportion of non-polar aliphatic LCRs, such as Gly-rich, Pro-rich and Ala-rich compositional biases, whereas proteins in the eCLIP-only set contain more polar LCR, especially LCRs containing the acidic Asp and Glu and positively charged Lys. It has been shown that such LCRs, especially those containing lysine, tend to condense into coacervates with RNA [38], which likely contributes to the RNA binding properties of these proteins. Interestingly, group 5, which yields the most unique motif signatures among groups in the eCLIP-only set, is most enriched for Ser bias compared to all other groups.

Finally, the value of PEKA-scores becomes clear when comparing them to the recall of groups in the *in vitro* set. The median PEKA-score of each group follows the same order as median recall: the highest in group 1, followed by 4, 3 and 2. Thus, the extent of motif enrichment (as quantified by PEKA-score) is closely related to the extent of cross-validation with *in vitro* data (Figure 6a,b). Taken together, the similarity index and PEKA-score appear to be valuable measures of the RNA sequence specificity of the studied proteins, as well as of data specificity.

## Discussion

Here we present PEKA, a software that examines individual CLIP data to reliably discover the motifs that reflect the specificity of purified RBPs. PEKA implements an approach to motif enrichment that effectively corrects the technical biases at crosslink sites by using the low-count crosslink sites as a source of background sequences. We use PEKA to move beyond summarised meta-motifs and gain insights into motif classes, their distribution patterns around crosslink sites, and their commonalities across datasets and across RNA regions. By performing comparative analyses of motif enrichments, we find that the inter-data specificity of enriched motifs, and the extent of motif enrichment (PEKA-score), are valuable measures of data specificity that relates to the domain composition of the studied proteins and their regional binding preferences. We provide a web platform for visualisations of these motif enrichments and distributions for motif-centric or RBP-centric data exploration of the entire ENCODE resource [4, 5].

The eCLIP experimental protocol omits the visualisation of purified protein-RNA complexes, which is normally used for experimental optimisation of specificity via visual analysis of expected vs. co-purified RBPs [6], and therefore analysis of data specificity is particularly pertinent to eCLIP. Previous studies evaluated successful IP, library complexity, and the number of reproducible CLIP peaks as a signature of data quality [3], however these metrics serve mainly as measures of data sensitivity [39]. As a measure of data specificity, previous studies have reported the agreement of enriched motifs between eCLIP with RBNS [3], but have not used such analyses to compare various quantitative metrics of data specificity across datasets. We find that 5% of datasets contain the highest specificity characteristics of high agreement with orthogonal *in vitro* data and inter-eCLIP data specificity, whereas ∼40% of datasets have low inter-eCLIP specificity, lack orthogonal *in vitro* data, and in most cases show a predominance of G-rich motifs. Notably, such motifs are found enriched even in some eCLIP (but not iCLIP or PAR-CLIP) datasets from the highest quality datasets, such as TIA1 (Figure 2). It has been hypothesized that these motifs may be bound by co-purified RBPs [3], and our analyses indicate that this contamination affects mainly eCLIP, but not iCLIP and PAR-CLIP datasets of the same proteins. Interestingly, one earlier study reported enrichment of G-rich motifs in PAR-CLIP background [40], however, we did not detect these motifs commonly enriched in PAR-CLIP data examined in our study. Since we find that enrichment of G-rich motifs is seen in datasets with preferential intronic crosslinking, we speculate that G-rich background might result from contamination of chromatin and associated RBPs. Such contamination tends to emerge when cell extracts are too concentrated and viscous [41], and therefore the conditions of CLIP normally recommend sonication and DNase treatment of a well-diluted extract, and visualisation of purified protein-RNA complexes to confirm lack of contaminating signal [42].

It is expected that PEKA-score and the resulting top motifs will be influenced by a peak file of choice, but nevertheless we show that the kmers identified by PEKA in this study correlate well with the mCross k-mers that used a different peak calling approach. Nevertheless, both approaches have in common that they use internal background for normalisation. However, the SMInput filtering, which is meant to decrease the extent of extrinsic background in CLIP data, sometimes leads to major changes in identified motifs. Surprisingly, we found no evidence of this approach increasing data specificity in general, while it leads to a substantial loss of data sensitivity. In particular, this approach tends to decrease specificity of data for RBPs that bind to U-rich motifs. In conclusion, we find that SMInput filtering does not necessarily increase data specificity, while it greatly decreases data sensitivity and can remove foreground signal from eCLIP data.

We find that RBPs with highest motif enrichments (high mean PEKA-scores) and inter-eCLIP data specificity (low similarity score) tend to have several RRM and KH domains and a high proportion of aliphatic LCRs (Figure 6). While this informs on the extent that RBPs might be sequence-specific [43], it also informs on the specificity of eCLIP datasets. Datasets that agree most highly with *in vitro* data are expected to be most specific (i.e., group 1 in Figure 6), and these have the highest PEKA-scores and inter-eCLIP data specificity. Interestingly, the same RBPs are also most efficiently identified by RNA-crosslinking purification (eRIC), indicating that they crosslink well. Interestingly, all RBPs in group 1 predominantly bind to introns. Thus, it is at present somewhat unclear whether the major differences in motif specificity (similarity index) and enrichment (PEKA-score) across eCLIP datasets reflect differences in the extent of biological sequence specificity of studied RBPs, variable capacity of eCLIP to isolate the studied proteins without co-purified RBPs, regional differences in motif-driven RBP recruitment, or a mixture of these factors.

Our analysis of motif enrichments is focused on protein-coding genes because most eCLIP datasets are for RBPs that do not bind to noncoding RNAs (ncRNAs), and partly because the RBPs that do primarily bind to a few abundant ncRNAs [3] that are not sufficient for motif derivation. Moreover, abundant ncRNAs are highly structured and modified, and involve complex and multi-step RNP assembly mechanisms, and therefore they require integrative analysis of multiple types of data and structural modeling before the functional sequence motifs can be extracted [44, 45]. Nevertheless, a subset of eCLIP data is highly enriched in ncRNAs, and clustering of eCLIP by regional crosslinking profiles that include abundant ncRNAs and repetitive RNA elements does lead to smaller clusters [3] compared to the clusters defined by enriched motifs in our study (Figure 5). Therefore, it is important to note that the similarity index from our study only represents specificity of motifs enriched in protein-coding genes, rather than specificity of eCLIP data as a whole. In the future, it will be valuable to analyse enriched sequence and structural motifs in combination with the types of bound RNA to understand additional features that contribute to the specificity of CLIP datasets.

## Conclusions

PEKA software enables discovery of enriched motifs from CLIP data while minimising the biases of crosslinking and of other technical steps of CLIP. We show that PEKA discovers motifs with various patterns of enrichment at crosslink sites, including those that are broadly enriched around crosslink sites, which could contribute to the condensation-dependent assembly as shown recently for TDP-43 [22]. In the future, it will be interesting to further investigate the cases where differences are observed between the motifs detected by PEKA and *in vitro* data, which may be due to biological factors, such as biased sequence characteristics of genomic regions, multi-protein assembly characteristics that cannot be reproduced *in vitro*, requirement of RNA modifications and post-translational modifications of RBPs for *in vivo* binding patterns, and variable specificity of eCLIP or RBNS data. We anticipate that assessment of the extent of motif enrichment (PEKA-score) and inter-data similarity of enriched motifs (the similarity index) will become increasingly valuable as further CLIP datasets become available. This will help to better understand the specificity of CLIP data, as well as the molecular mechanisms that determine the specificity of protein-RNA interactions in cells.

## Methods

### Source data and primary analysis

Many variants of CLIP protocol enable identification of crosslink sites by analysis of crosslink-induced features. Here we follow the approach introduced by the iCLIP method, where cDNA truncations serve to identify crosslink sites, which has been adopted by many other variants, including eCLIP [2]. Due to the short time of excitation, ultraviolet light is considered a zero-distance crosslinker that induces the formation of a covalent bond between the RBP and RNA that are in direct contact at the time of irradiation [46]. While it is in principle possible to derive some crosslink sites also from analysis of mutations in eCLIP reads, we did not use this information as these sites were less effective with regards to specificity of derived motifs [14], and the proportion of reads with such mutations varies between eCLIP datasets in a way that could introduce further technical variation. Thus, to obtain crosslink sites, eCLIP fastq files were downloaded from the ENCODE consortium (Additional file 2) [4, 5] and processed to get positions of cDNA truncations [47]. First, adapters were removed with Cutadapt (v3.4) using two rounds of adapter removal to account for double ligations as per the ENCODE standard operating procedure and the unique molecular identifier positioned at the end of the fastq header. We aligned the second read as this contained information as to the crosslink position [28]. We used the nf-core/clipseq pipeline [48, 49] to process the reads, first filtering out reads that aligned to rRNA or tRNA and then aligning to the human genome (GRCh38 primary assembly, Gencode V29 annotation). PCR duplicates were removed using unique molecular identifiers and the crosslink position identified as the coordinate immediately 5’ to the alignment start. eCLIP narrowPeaks, combined from both replicates and filtered with corresponding SMInput control, were also downloaded from the ENCODE consortium [4, 5].

For iCLIP samples for hnRNPC [50], TIAL1 [51] and TARDBP [52], bed files with positions of cDNA truncations were obtained by standardised iCLIP read processing employing the iMaps web server [19] mapping to the human GRCh38 genome build. For PAR-CLIP samples for TIA1 [53]; IGF2BP1, IGF2BP2, PUM2 [54]; FUS, EWSR1, TAF15 [55]; and hnRNPC [56], fastq files were downloaded from the SRA database. Reads were stripped of the 3’adapter sequence by Flexbar (v2.5) and collapsed to remove PCR duplicates. Next, reads were sequentially mapped to reference transcripts by Bowtie2 (v2.3.2) in the following order by retaining the unmapped reads from the previous to the next mapping step. We started with human pre-rRNA (GenBank U13369.1), followed by rRNA (GenBank NR_023363.1, NR_003285.2, NR_003287.2, NR_003286.2), snRNA, snoRNA, other ncRNA (all from Ensembl, including RN7SL), tRNA (GtRNADb), mtDNA (GenBank AF347015.1) and finally the human genome (GRCh38, primary assembly). The last genome-mapping step was performed by the STAR aligner (v2.5.3a) and only uniquely mapped reads (MAPQ=255) were retained for further processing. T-C transitions were extracted using the SAMtools mpileup command and row_mpile_coverage_plus_TC.pl script [57]. SRA accession codes to all iCLIP and PAR-CLIP samples used for this study are available in Additional file 3.

We assigned crosslink sites by positions of cDNA truncations for eCLIP and iCLIP data (located 1nt upstream of 5’ mapped read positions), and by positions of T-C transitions for PAR-CLIP data. We didn’t use mutations from eCLIP and iCLIP experiments, because mutations were found to be less consistent across replicates for motif discovery [14], the proportion of mutations can vary greatly across experiments, and the causes of mutations can also derive from sequencing errors or divergence from genomic reference sequence, among others. Crosslink files of replicate experiments for each RBPs from each study were then merged, such that if multiple replicates identified the same crosslink site, cDNA counts were summed up (as defined in Additional file 3).

All remaining analyses of CLIP data were done using newly-developed or modified scripts described below, written in Python 3.7.3., using GRCh38 genome build and GENCODE version 30 annotation, which was used to produce segmentation file with iCount segment.

5-mer z-scores for the RNA-Bind-n-Seq datasets were kindly provided by Dominguez laboratory [15]. For each dataset, the z-score values for the concentration with the highest enrichment were used. RNAcompete 7-mer z-scores were obtained from [[58]] web supplementary data. Raw and normalized mCross 7-mer z-scores for all ENCODE eCLIP datasets were provided by Zhang laboratory [14]. 7-mer z-scores were converted to 5-mer enrichment scores by calculating the arithmetic mean of all 7-mer scores that contain a given 5-mer. For the ease of reproducing findings of our study, we provide the 5-mer z-scores for RBNS, RNAC and mCross datasets in Additional file 4 and 5-mer rankings derived from these z-scores in Additional file 6, but alert the readers that all of these originate from the previous studies referenced above.

Enhanced RNA interactome capture (eRIC) data from Jurkat cells was obtained from [Supplementary Data 1 in [36]]. For our analyses, we visualised the log2-fold change (FC) in signal intensity in UV irradiated (UV+) over non-irradiated (UV-) eRIC samples.

### Peak-calling

For the purpose of motif analysis, peaks of crosslinking events were produced with the Paraclu script [21] the Paraclu peak calling is very fast. Since Paraclu doesn’t use any information of genomic annotation, we wanted to avoid that peaks would arise simply due to increased coverage of crosslinks within exons. For this purpose, we ran Paraclu twice, first on the whole genome and then on just the exonic portion of the genome, using the segmentation file produced from gencode hg38 v30 annotation with iCount segment. Crosslinks that lie in exonic regions of the same genes were combined prior to peak calling to mimic the spliced transcript, and a 1kb spacer was inserted between genes to avoid peaks being called across genes. For running Paraclu on exonic peaks, the parameters were set to minValue of 10 and max cluster length of 100nt, while for genomic peaks, the parameters were minValue of 10 and max cluster length of 200nt. Peaks overlapping by >75% with exons were removed from genomic peaks, and then these filtered genomic peaks were merged with exonic peaks.

### Positionally Enriched k-mer Analysis (PEKA)

PEKA is a tool for finding enriched sequence motifs from CLIP data. We focused on analysis of crosslink sites in the protein-coding regions. First, thresholding splits the crosslinks into high-confidence thresholded crosslinks (tXn) and reference background crosslinks (oXn). The thresholded crosslink sites (tXn) are indicative of high-occupancy protein binding, and the reference group (oXn) can be referred to as ‘intrinsic background’ of CLIP experiments, stemming from more transient RNA binding of the protein [6].

The first step is to use regional thresholding to obtain tXn. Each transcript is considered separately, such that all exons (including CDS, UTRs and ncRNAs) within a gene are combined into one region, and each intron and intergenic region are treated as their own region. Within each region, a cDNA count threshold is determined at which ≥70% (the default of the code, but this percentile can be modified by the user) of the crosslink sites within the region have a cDNA count equal or below the threshold, e.g. if the region contains 10 crosslinks, nine out of which have a cDNA count of 1 and one has a cDNA count of 2, the threshold for that particular region is set to 1. tXn are then identified that have cDNA count above the threshold, and overlap with the peaks that are provided by the user. On the other hand, oXn are all crosslinks that fall outside of peaks. If the number of crosslinks is very abundant, tXn and oXn are randomly sampled to obtain 1 million tXn and 3 million oXn positions, which was found to yield comparable data in all tested conditions, and can save memory and reduce the time of computation. While this sampling is done by the default, it can be turned off by the user.

By default, PEKA examines motif-enrichment in the following transcriptomic regions: introns (from both coding and non-coding genes); 3 ′UTRs; other protein-coding exon regions (comprised of coding-sequence exons and 5 ′UTR), non-coding RNAs (comprised of exons from non-coding RNAs); intergenic regions; protein-coding genes (comprised of full sequence of protein-coding genes); and whole genome. Optionally, PEKA also allows separate analysis of 5 ′UTRs and coding-sequence exons. PEKA also supports the use of repeat-masked genomes, and provides the options to either fully exclude repeat elements from motif-enrichment analysis, to plot the enrichment in repeat-masked genome in uppercase letters, and that within the repeats in lowercase letters, or to analyze motif-enrichment exclusively in repeating regions.

Next, foreground and background sequences are extracted around tXn and oXn, respectively (Figure 1 b), and used to conduct enrichment analysis separately in each transcriptomic region. Foreground sequences are genomic regions spanning −150…150nt around tXn and are subdivided into the proximal (−20…20nt around tXn) and distal windows (−150…−100 and 100…150 nt around tXn). Background sequences consist only of the proximal region and span −20…20nt around oXn. The values used for proximal and distal regions used in our study are set by default in the code, but can be adjusted by the user. However, we recommend that the selected proximal region is not more than −50…50nt, as we rarely see enrichment of relevant motifs further than 50nt from crosslink sites.

For each k-mer, PEKA scans across the foreground and background sequences and records the presence of a k-mer by assigning count 1 if present or 0 if absent (Figure 1c); for k-mers of odd lengths, the position that coincides with the middle of the k-mer is assigned the count and for k-mers of even lengths, the position that corresponds to its length divided by a factor of 2 is assigned the count (i.e., 3rd overlapping nucleotide for a 6mer). For sequences located in each genomic region, mean k-mer counts are calculated at each position to obtain k-mer occurrences around crosslink sites. Afterwards, relative k-mer occurrence is calculated by dividing k-mer occurrence at each position with the mean k-mer occurrence across all positions within distal windows of the foreground sequences. Relative k-mer occurrence is calculated separately for the foreground sequences (to get RtXn) and for 100 samples of randomly selected background sequences of the same number as the number of foreground sequences (to get RoXn).

PEKA calculates enrichment of each kmer by analysing the positions (relative to crosslinks) where kmer is present above a background level, i.e., ‘the relevant sequence positions’ (Figure 1d). These positions are identified by analysis of the relative occurrences, such that each sequence position within the proximal window gets its own position-threshold value, calculated from the 100 RoXn distributions. The position-threshold is defined based on all RoXn values at that position across all possible k-mers (100*4^k^ values), and is equal to the value at which a specified percentile of evaluated RoXn values at that position fall below the threshold. This percentile is set to 80% for 5-mers and for each k-mer, the positions that contain RtXn higher than the position-threshold value are marked as relevant.

Finally, PEKA-score is calculated by analysis of k-mer values at the relevant sequence positions (Figure 1f). For each k-mer, a mean RtXn (ARtXn, i.e. approximated occurrence around tXn) and 100 mean RoXns (ARoXns, i.e. approximated occurrence around oXn) are calculated across the relevant positions, and PEKA-score is calculated as (ARtXn - mean(ARoXn)) / std(ARoXn), where mean(ARoXn) represents the mean of ARoXn values across all 100 samples, and std(ARoXn) represents the standard deviation of ARoXn values across all 100 samples (Figure 1f). The k-mers are then ranked by PEKA-score from the most to the least enriched and all results are given in a table. PEKA-scores for all datasets analyzed in the scope of this research are available in Additional file 5 and 5-mer rankings derived from these scores are available in Additional file 6.

To aid user in the first stages of analysis, PEKA outputs visualisation of occurrence profiles based on the occurrence of top 20 (or as many as defined by the user) k-mers is visualised in graphs that group together the motifs with similar sequence characteristics and occurrence distribution (Suppl. figure 1, Additional file 1). Sequence similarity is stored as a matrix of pairwise jaccard indices calculated with the python *textdistance* module [59]. The similarity of k-mer occurrence distribution is measured with 1) Spearman rank correlation coefficient, 2) occurrence maximal values and 3) max peak position. Matrices containing sequence-based and distribution-based metrics are combined at varying weights, the resulting matrix is used for clustering, and the optimal clusters are selected for visualisation based on the lowest standard deviation of occurrence medians within clusters.

### Sequence based clustering of k-mer groups

For sequence-based clustering of k-mers, individual motifs are first converted into tokens that reflect their sequence properties. Tokens resulting from a k-mer are all its subsequences, each combined with an end number denoting their incidence within a k-mer. For example a k-mer ‘AGGU’ is tokenized into: ‘A1’, ‘AG1’, ‘AGG1’, ‘AGGU1’, ‘G1’, ‘GG1’, ‘GGU1’, ‘G2’, ‘GU1’, ‘U1’. Notice that two guanosines in the example k-mer produce two distinct tokens ‘G1’ and ‘G2’, due to the addition of token occurrence at the end. After tokenization, pairwise Jaccard similarity is calculated for all k-mers in the group. Jaccard similarity is a quotient of the number of shared tokens between two compared motifs and the number of all tokens in the union formed by the k-mers. The k-mers are then clustered with an affinity propagation method, based on the resulting similarity matrix. Affinity propagation clustering was implemented with the scikit-learn v0.21 python module, using the damping parameter of 0.5, the maximum allowed number of iterations set to 1000, and the number of convergent iterations set to 200. Affinity propagation clustering automatically determines the number of resulting clusters.

For Suppl. figure 2 (Additional file 1), the similarity matrix was obtained for the top 20 k-mers as described and converted to the distance matrix (1-similarity matrix) for K-means clustering (implemented with the scikit-learn v0.21 python module) to split the motifs into two clusters.

### k-mer logos and consensus sequences of PEKA k-mer groups

To visually represent each k-mer consensus groups, obtained after sequence-based clustering, we used sequence logo representations. These were created by k-mer multiple-sequence alignment (MSA) transformed to position-frequency matrix (PFM), as follows. First, pairwise sequence alignments of k-mers are obtained by employing global Needleman-Wunsch algorithm with the *skbio.alignment* module [60], version 0.5.1, setting the score for a nt match to 2 and the mismatch score to −1. Other scoring parameters are left on their default settings - penalty for opening the gap is set to 5, the penalty for gap extension is 2 and terminal gaps in alignment are not penalized. Then, pairwise alignments are collated into the MSA, starting with the highest scoring alignment (in case there are multiple alignments with the same score, the one that is the first by alphabetical sorting is taken) and aligning the second best pairwise alignment containing one of the motifs already included in the MSA to it. This process is repeated with the next best scoring pairwise alignment, until all kmers are aligned in the MSA. MSA is then transformed into a PFM, which denotes the frequency of each nucleotide at each position within the alignment. PFM is used to plot sequence logos with the *seqlogo* [61] module, version 0.1.9. By using a rolling window of a predefined length, a motif consensus can also be determined from the PFM by sliding the window across all PFM positions and summing the occurrences of nucleotides within a window. Where the sum is greatest, the majority consensus sequence is derived from the PFM. In case of ties between two or more nucleotides, IUPAC nucleotide notation is used. In case of multiple windows with the same highest scoring sum, the first window in the PFM to get that score is used to derive the consensus sequence. It should be noted that k-mer logos are only an approximate and not an accurate representation of the binding motifs, as PFMs are generated solely based on the sequence alignment of k-mers. Thus, k-mer logos do not necessarily reflect the precise relative positioning of the k-mers or their frequency in the foreground sequences that were used to identify the k-mers. Rather, k-mer logos are used to aid in the visualisation of the common sequence features of each investigated group of k-mers.

### Clustering eCLIP datasets

For Figure 5 we clustered eCLIP datasets based on their distribution within the regions comprising protein coding genes (namely 5 ′UTR, CDS, 3 ′UTR and intron), as well as on their 5-mer ranks. We first calculated cosine similarities between datasets for the regional information and 5-mer ranks separately, and then added the similarity matrices with weights 0.3 and 0.7, respectively. Then, we transformed the combined cosine similarities into cosine distances and used these to perform hierarchical clustering with the *scipy.hierarchy* module.

For Figure 6 we first split the eCLIP datasets into two groups based on whether or not they had available orthogonal *in vitro* data and then performed K-Means clustering (implemented with the scikit-learn v0.21 python module) on each group, using either an equally weighted combination of similarity index and recall, or just similarity index where no *in vitro* data was available. Prior to clustering, we normalized recall and similarity index across eCLIP datasets using min-max scaling, to ensure an equal contribution of these parameters to clustering. eCLIP datasets with available *in vitro* data were split into 4 clusters and datasets without available *in vitro* data were split into 3 clusters. For heatmap visualisation, we arranged clusters in each group by their median similarity index, and additionally, datasets within each cluster were arranged in ascending order based on their similarity index. eCLIP clusters and data related to the main heatmap in Figure 6a are available in Additional file 7.

### Recall of PEKA motifs against *in vitro* data

Recall was calculated from a confusion matrix, using the ranking of motifs from *in vitro* datasets (RBNS or RNAC) as ground truth towards which the ranking of PEKA motifs for orthogonal CLIP datasets were compared. In both datasets, k-mers were classified into two groups, based on their ranking: 1) binders and 2) non-binders. The rank cutoff for binders was set to 20 for *in vitro* datasets and to 50 for CLIP datasets. Thus, recall in this study represents a proportion of top 20 motifs from *in vitro* dataset that are found among the top 50 motifs in the corresponding CLIP dataset. Recall values for all eCLIP datasets are available in Additional file 7.

In Figures 3, 5, 6, and Suppl. Figure 2 (Additional file 1), we compared selected eCLIP data (Additional file 2) with RBNS or RNAC. If both *in vitro* datasets were available for a particular protein, we always prioritized RBNS over RNAC for recall calculation as RBNS z-scores were readily available for 5-mers, whereas RNAC required transformation from 7-mer to 5-mer scores.

### Similarity score

Similarity score compares how similar the top motifs of a particular eCLIP dataset are relative to other datasets. Similarity score was obtained by calculating pairwise overlap ratios on top 50 kmers for all eCLIP datasets and then calculating the mean of these overlap values for each dataset. Similarity score of 0 would indicate that the top 50 of k-mers in a certain dataset were not ranked among the top 50 in any other dataset. In contrast, higher values of similarity score indicate that top motifs of a specific dataset overlap with top motifs in many other datasets. Similarity scores for all eCLIP datasets are available in Additional file 7.

### Metaprofile of average motif coverage around crosslinks

This script plots metaprofiles of average motif coverage, and is available at github [62]. It visualises the average k-mer coverage of a motif-group around crosslink sites within specified transcriptomic regions (corresponding to regions described in PEKA). By default (and for this study), the analysis is performed on the full set of user-provided crosslink sites by considering their cDNA counts, but optionally these sites can be filtered to determine thresholded crosslinks (tXn, as described above). Additionally by default (and for this study), cDNA count of crosslink positions that were identified by more than 20 unique cDNAs was capped at 20 to avoid the most abundant RNAs from dominating the signal, but this can be modified by the user.

Sequences flanking the crosslink sites (default window is −150…150 around the crosslink site, but it can be modified by the user) are scanned with a rolling window equal to k-mer length to find parts of the sequence that match k-mers from the investigated motif group. All positions containing a motif from the investigated group are given a score corresponding to the cDNA count of the evaluated crosslink position and the remaining positions are scored 0. Scores at each position around crosslinks in the assessed region are summed and divided by the total cDNA count of all evaluated crosslinks to generate the coverage showing the percent crosslink events overlapping with any k-mer from the group at each position. Optionally, the user can select to visualise the coverage unweighted by cDNA count, in which case all positions containing a motif from the investigated group are given a score of 1, remaining positions are scored 0; scores at each position around crosslinks in the assessed region are summed and divided by the number of evaluated crosslink sites to get the coverage.

Finally, coverage distributions smoothed using rollmean function with window size of 6 (can be modified by the user) and the metaprofiles for the list of analysed crosslink files, i.e. samples, are plotted on the same graph. For analysis in Figure 5, we calculated the average motif-group coverage weighted by cDNA scores around tXn and oXn in the ‘whole gene’ region for 24 eCLIP datasets.

For the web interface (see [20]), we calculated metaprofiles of average motif coverage across 24 motif groups encompassing all 5-mers, for all eCLIP datasets. For input, we used thresholded crosslink sites in the ‘whole gene’ region, obtained by PEKA, either removing crosslinks from genomic repeats (no repeats) or not (with repeats). We calculated the weighted motif coverage for each motif group in each sample for ‘intron’, ‘other exon’, ‘3 ′UTR’ and the ‘whole gene’ region. For each region, we visualised the coverage profiles for 40 datasets with the highest maximal coverage value within the window −50…50 around crosslink sites in heatmap format, clustering the datasets based on the metaprofile similarities and arranging the datasets within each cluster by falling max coverage values.

For the web interface we also created regional scatterplots for each motif-group (see an example at [63] displaying on the y-axis the maximal value of motif-group coverage for each dataset in the window −50…50 around crosslink sites for the analyzed region, and on the x-axis the maximal enrichment in the selected dataset compared to all datasets at the same position (z-score). Z-score value was obtained at each position within the window −50…50 around crosslink site by calculating the difference between the coverage of the investigated dataset and the mean coverage across all datasets at the same position, and dividing the resulting value with a standard deviation of coverage values at specified position. The scatterplots display a maximal z-score achieved by each dataset.

### Generation and selection of motif groups and RBPs for visualisation

The differentially enriched motif groups for Figure 4 were generated as follows. We obtained k-mers that were differentially enriched between *in vitro* approaches (RBNS or RNAC) and eCLIP motifs identified by PEKA, mCross or the ‘local’ approach (RtXn) for all RBPs where data were available for at least one *in vitro* methods and for mCross. For each dataset, we calculated differences in k-mer ranks by subtracting the ranks in the method used to analyze eCLIP datasets from their respective *in vitro* ranks. Then, a mean difference was obtained for each k-mer across all eCLIP datasets and a z-score was calculated for each mean difference. By applying a confidence interval of 95% (z-score < −1.96 or > 1.96), we obtained significantly enriched and depleted k-mers for each approach.

### Analysis of domain types and compositional biases

We downloaded gff files from UniProt for all analyzed RBPs and filtered the features for Domains and Compositional bias. To decrease the number of bias types for the sake of presentation, names of compositional biases were simplified to only contain the code for the dominant amino acid. For example Gly-rich and Poly-Gly biases were changed to ‘Gly’. If a specific domain or compositional bias occurred less than 5 times across all datasets, we marked it as ‘other’. This was then presented in Figure 6 by counting the number of each domain type and compositional bias within each cluster of datasets, and normalizing the counts with the number of RBPs within the cluster to generate stacked bar plots. Number of compositional biases and KH/RRM domains are available for all eCLIP proteins in Additional file 7.

## Supplementary information

### Additional files

**Additional file 1** contains Supplementary Figures 1-4.

**Additional file 2** contains accession codes and metadata for ENCODE eCLIP datasets used in this study and specifies eCLIP datasets that were used to generate the Figures 1-6 and Suppl. Figures 1-3 (Additional file 1) in this manuscript.

**Additional file 3** lists iCLIP and PAR-CLIP experiments related to Figure 2.

**Additional file 4** lists 5-mer z-scores for RBNS, RNAC and eCLIP datasets analyzed with mCross, which were obtained as described in Methods.

**Additional file 5** lists 5-mer PEKA-scores for eCLIP datasets processed with Paraclu peaks and narrowPeaks, iCLIP and PAR-CLIP datasets.

**Additional file 6** lists 5-mer ranks for all datasets used in this study, i.e. eCLIP datasets analyzed with PEKA (using Paraclu peaks or narrowPeaks), eCLIP datasets analyzed with mCross, as well as iCLIP, PAR-CLIP, RBNS and RNAC datasets. For RBNS, RNAC and mCross, additional data are available, which are not shown in the Figures.

**Additional file 7** includes data related to Figure 6 of this manuscript, namely similarity index, recall, number of KH/RRM domains, number of LCR, log2FC eRIC, and mean PEKA-score across top 50 k-mers.

### Supplementary figure legends

**Supplementary Figure 1 | Positional motif clusters of top 20 ranked motifs for TIA1 eCLIP in HepG2 cell line.**

After ranking k-mers based on their PEKA-score, top n k-mers are clustered, based on their sequence and their occurrence distribution around tXn. Occurrence profiles for k-mers within each cluster are plotted together on a graph and each cluster is represented by its consensus sequence. The final plot shows summed k-mer occurrences for each cluster to elucidate which k-mer groups dominate the RBPs binding landscape and what is the group’s approximate distribution pattern.

**Supplementary Figure 2 | Comparison of PEKA and mCross motif analysis.**

**a)** Comparison of PEKA and raw mCross in their ability to recover the top 20 motifs from *in vitro* dataset in corresponding eCLIP data. The lines show a mean percentage of top 20 *in vitro* 5-mers that were recovered among the top n motifs in each method, across 41 eCLIP datasets for which RBNS or RNAC data was available (representing 28 distinct RBPs in total). Shaded area represents the standard deviation across evaluated datasets at a certain threshold of top PEKA/mCross k-mers. **b)** k-mer logos for two clusters of top 20 5-mers (see Methods) obtained for TIA1 in HepG2 cell line by PEKA, normalized mCross and raw mCross. All three approaches to motif analysis retrieved the canonical U-rich TIA1 motifs. PEKA and raw mCross also found the G-rich motifs enriched among the top 20 5-mers, but these were removed by normalization process, performed by mCross. **c)** Sequence logos for TIA1 eCLIP in HepG2 cell line downloaded from the mCross base [14]. The sequence logos were derived from the top 10 7-mers in the dataset, ranked by normalized mCross z-scores. 7-mers were clustered to produce sequence logos with Stamp [64] as described in [14].

**Supplementary Figure 3 | Clusters of top k-mers enriched in hnRNP C eCLIP and in corresponding SMInput**

**a)** Occurrence profiles of top 18 k-mers for hnRNP C eCLIP in HepG2 cell line, grouped into 4 clusters by PEKA. All four clusters contain U-rich motifs. **b)** Metaplot of summed k-mer occurrences for each cluster for hnRNP C eCLIP. **c)** Occurrence profiles of top 20 k-mers for hnRNP C SMInput in HepG2 cell line, grouped into 3 clusters by PEKA. The first cluster contains U-rich motifs, the second is dominated by C-rich motifs and the third by C/U-rich motifs. **d)** Metaplot of summed k-mer occurrences for each cluster for hnRNP C SMInput.

**Supplementary Figure 4 | k-mer logos for groups of differentially ranked 5-mers between RBNS and eCLIP.**

k-mer logos show characteristics of 5-mers that were differentially ranked between RBNS and eCLIP, analyzed by PEKA, mCross or ‘local’ approach. k-mer groups that were differentially enriched in eCLIP, relative to RBNS, are on the left and groups of k-mers that were depleted in eCLIP, relative to RBNS, are shown on the right. k-mer logos are generated from aligned k-mers within each group and thus represent general k-mer features in each group.

## Declarations

### Ethics approval and consent to participate

Not applicable.

### Consent for publication

Not applicable.

### Availability of data and materials

The source code for PEKA software is available at github [18], and is incorporated as part of the iMaps platform [19], where it can be used for interactive and password-protected analysis of uploaded CLIP data. Code for processing the ENCODE eCLIP data is available at ulelab/peka-eclip github repository [47] and the nf-core/clipseq pipeline is available at [49]. The source code for plotting metaprofiles of average motif coverage is available at [[62]].

Fastq files and narrowPeaks for all eCLIP datasets and SMInput controls analyzed in this manuscript are available from the ENCODE consortium [4, 5], with corresponding accession codes and download links listed in Additional file 2. PAR-CLIP data analyzed in this manuscript were obtained from various sources and can be downloaded from GEO or the SRA database (Additional file 3). iCLIP data analyzed in this study was obtained from various sources and bed files containing read starts can be downloaded from the iMaps v2 web server (Additional file 3) [19], and also from GEO or Arrayexpress databases. 5-mer z-scores for RNA-Bind-n-Seq datasets were provided by Dominguez laboratory [15], RNAcompete 7-mer z-scores were obtained from [[58]] web supplementary data, and raw and normalized mCross 7-mer z-scores were provided by Zhang laboratory [14]. 5-mer z-scores derived from original datasets as described in Methods are available in Additional file 4, and corresponding 5-mer ranks are available in Additional file 6. 5-mer PEKA-scores and corresponding ranks for all CLIP datasets represented in this manuscript are available in Additional file 5 and Additional file 6, respectively.

Data related to Figure 6 of this manuscript, namely similarity index, recall, number of KH/RRM domains, number of LCR, log2FC eRIC, and mean PEKA-score across top 50 k-mers, are available in Additional file 7.

Enhanced RNA interactome capture (eRIC) data from Jurkat cells is available from [[36], Supplementary Data 1].

### Authors’ contributions

AGA wrote the source codes for PEKA software and the software for plotting metaprofiles of average motif coverage. KK performed the computational analyses and produced the figures and tables for the manuscript and for the web platform. KK and JU wrote the manuscript. All authors read and approved the final manuscript.

## Acknowledgements

We thank Anob Chakrabarti for processing of eCLIP data and Charlotte Capitanchik for her help with the development and testing of the PEKA software, and Nina Lenaršič and Anja Trupej for their help with figure editing. We also thank Anob Chakrabarti, Charlotte Capitanchik, Ira Alexandra Iosub, Miha Milek and Miha Modic for their detailed feedback and guidance while editing the manuscript. We also thank prof. Daniel Dominguez and prof. Chaolin Zhang for kindly providing 5-mer z-scores for RNA-Bind-N-Seq and 7-mer z-scores for the mCross datasets, respectively.

