## Supplementary figures and images for "Positional motif analysis reveals the extent of specificity of protein-RNA interactions observed by CLIP"

### Additional file 1 - Supplementary Figures 1-4

Suppl. Figure 1

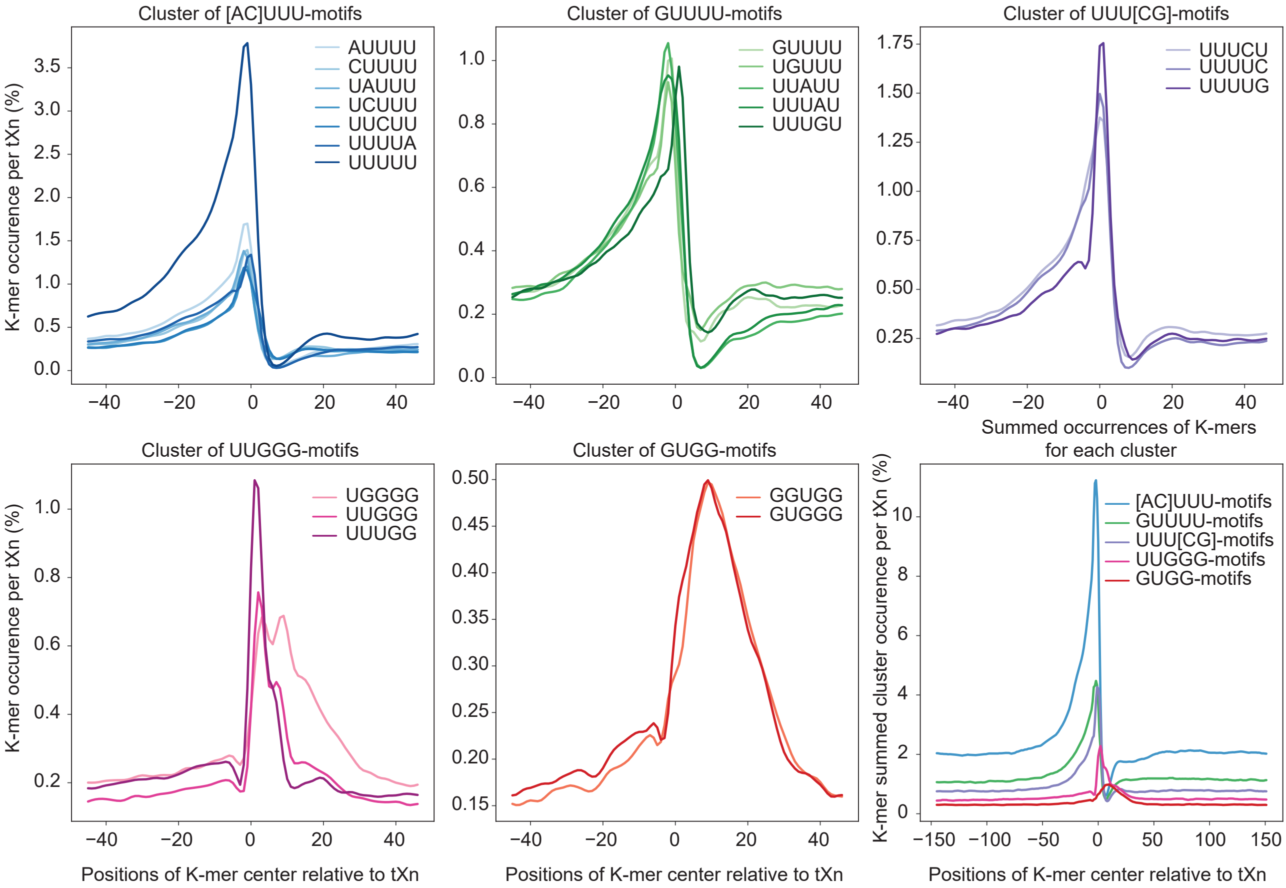

Suppl. Figure 2

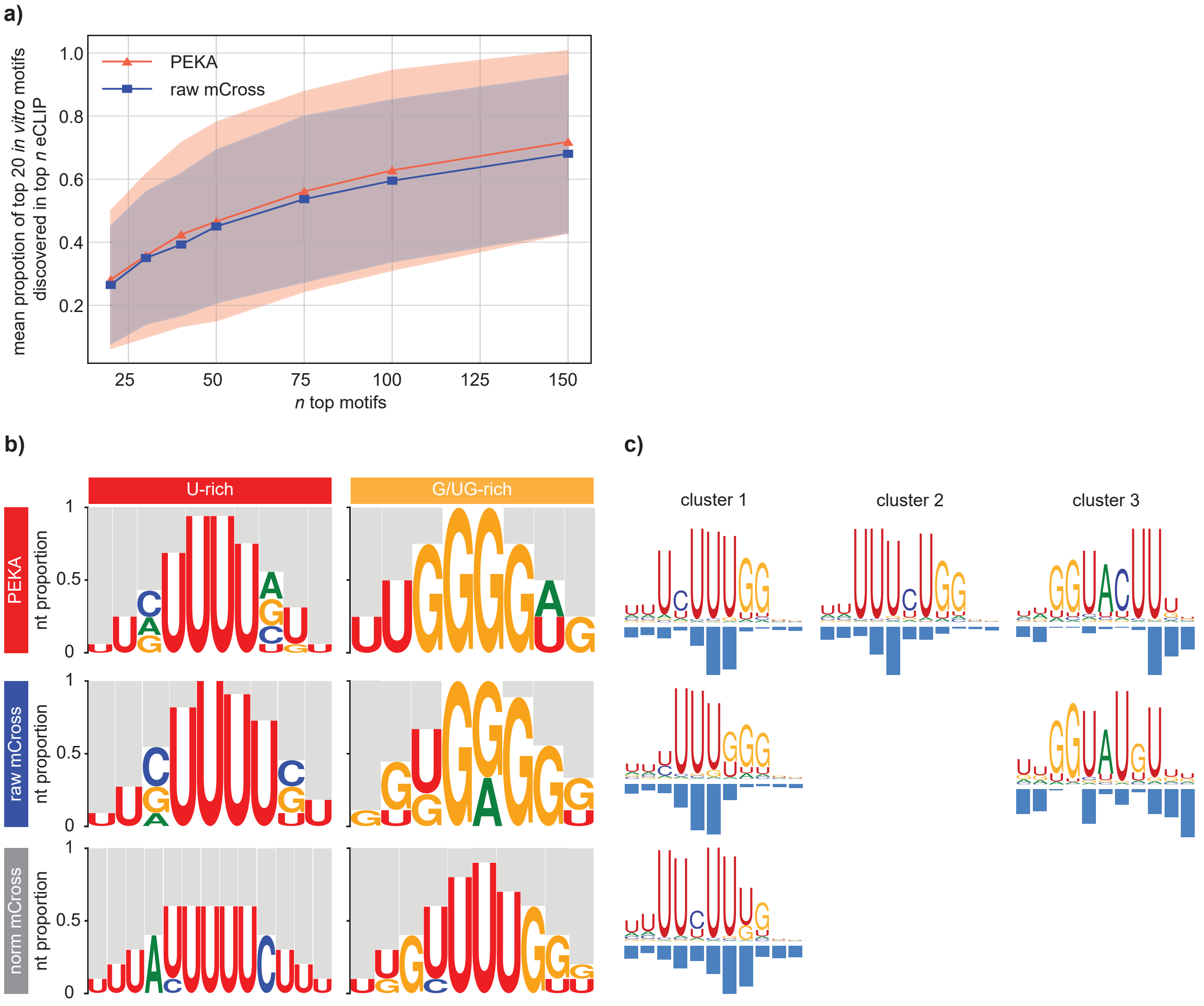

Suppl. Figure 3

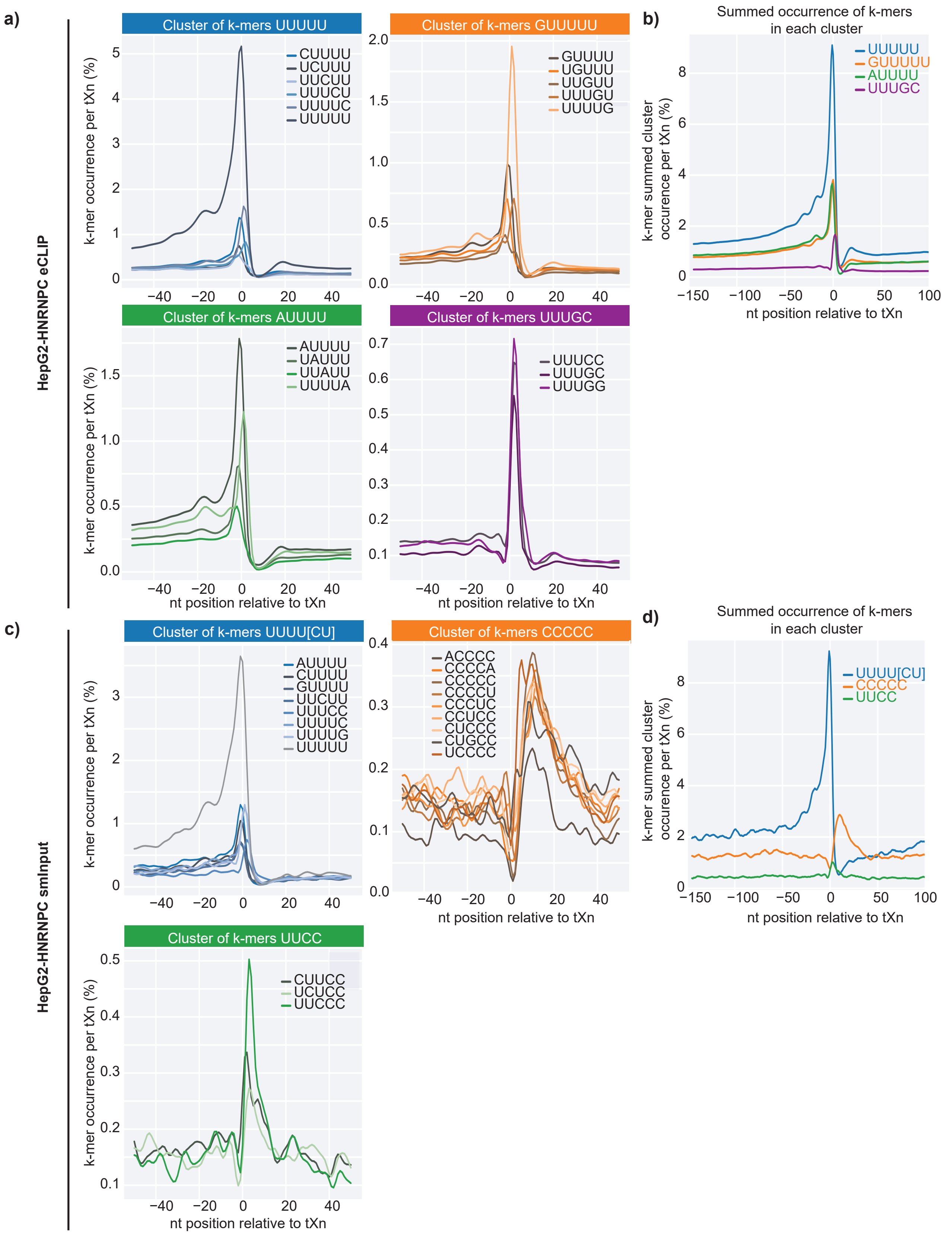

Suppl. Figure 4

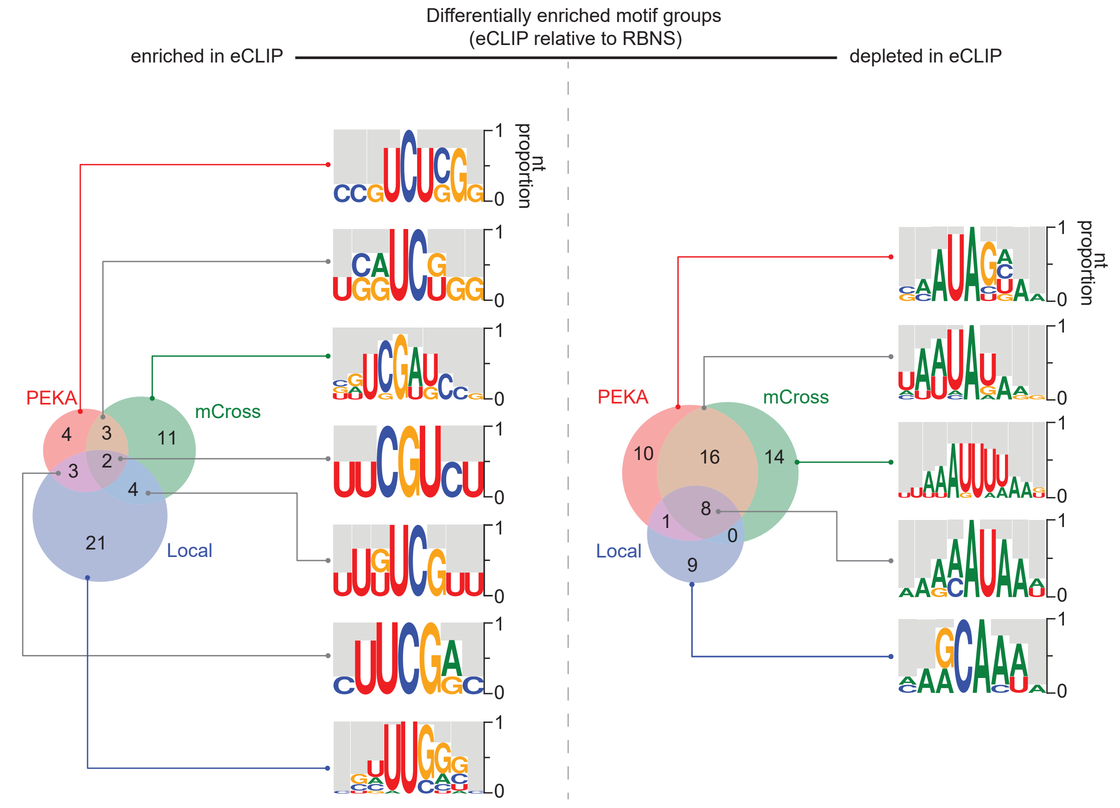
